# The Demographic and GDP Impacts of Slowing Biological Aging

**DOI:** 10.64898/2026.01.22.701157

**Authors:** Raiany Romanni-Klein, Nathaniel Hendrix, Richard W. Evans, Jason DeBacker

**Affiliations:** Harvard Medical School; Stanford University; University of Southern Carolina; Abundance Institute

**Keywords:** Aging, economics, demographics, labor supply, productivity, macroeconomics

## Abstract

Biological aging imposes significant socio-economic costs, increasing health expenses, reducing productivity, stalling population growth and straining social systems, culminating in reduced economic activity. We draw insights from interviews with 102 scientists working on aging biology and develop four macroeconomic simulations: slowing brain aging, slowing reproductive aging, and an overall delay in biological aging (including the novel concept of “replacing aging”). Our model is calibrated to represent how slowing biological aging manifests in the US economy and population through the channels of mortality, fertility, and productivity rates by age. We simulate the economic and demographic impacts of near-future advancements in aging science. We find that a one-year delay in brain aging alone could add $201 billion annually to US GDP. A one-year delay in overall biological aging could boost GDP by $408 billion annually, yielding $27.1 trillion in net present value in the long run.

## Introduction

The world is aging. By the mid 2030’s, people aged 80 and older will outnumber infants.^1,2,3^ More births would be a long-run solution to the emerging shortage of working-age adults worldwide. Yet higher fertility rates alone would be no silver bullet. In the short term, they worsen dependency ratios (newborns don’t work, and they temporarily remove their parents from the workforce); in the long term, they do not change the socio-economic costs of aging. By 2029, the United States will spend $3 trillion yearly — half its federal budget — on the medical treatment and social care of adults aged 65 or older.^4^ By 2040, China will be home to nearly half a billion people over 60.^5^ By 2050, Japan’s population will shrink by 20 million, while close to half its remaining population will consist of dependents, most of them retirees.^6^

Biological aging imposes significant socio-economic costs, increasing health expenses, reducing productivity, stalling population growth and output, and straining social systems. Because of this high dimensionality, any single valuation of breakthroughs in aging science is incomplete. In this article, we propose a macroeconomic approach to valuing breakthroughs in aging biology. We apply this method to simulate the value of four counterfactual advancements in aging biology. The advantages of our approach to valuing breakthroughs in aging biology include (i) the broad, macroeconomic nature of our measure, (ii) its natural monetary value, (iii) its general equilibrium nature, which incorporates the interactions of individuals, businesses, and policy in the economy, and (iv) its short- and long-run outlook, which considers population dynamics and lags in effectiveness and phase-in time.

Most previous economic valuations of medical breakthroughs are based on value of statistical life (VSL) and quality adjusted life year (QALY) methodologies. These approaches typically rely on surveys of individuals, natural experiments that reveal the trade-offs people make between risk and money, and VSL-calibrated partial equilibrium life cycle models of an individual life.^7,8,9,10,11^ Our macroeconomic model includes many individual and macroeconomic outputs, but we focus on its measure of the effects on gross domestic product (GDP). GDP provides a measure more directly related to changes in income, consumption, and tax revenue—key metrics considered by businesses, individuals, and governments. VSL and QALYs, by contrast, perceive life and health as valuable in their own right. We acknowledge this feature of the existing literature is extremely important. Our aim with this paper has been to supplement existing approaches by measuring some of the more direct economic impacts of advancements in aging science.

To our knowledge, ours is the first study to use a macroeconomic model and its GDP output to measure the value of advancements in aging biology. This macroeconomic approach has four major strengths. First, a measure of GDP generated from a general equilibrium model covers a broad range of individual incentives, responses, interactions, and feedback. For instance, our model captures how slowing reproductive aging might increase fertility rates, in turn increasing labor supply decades in the future and eventually leading to marginally downward pressures on hourly wages. This broad characteristic captures the multidimensional nature of the benefits to both individuals and the entire population of reducing the effects of aging.

A second strength of using GDP is its native monetary value. GDP is valued in the currency of the macroeconomic model. As such, no conversion is required from theoretical units of individual welfare or utility. This means our results measure the possible *direct* impacts on the economy of advancements in aging science. A third strength is more nuanced. The general equilibrium environment of the model we use captures feedback from individual choices, resulting population dynamics, prices, and the effects of government policies. This is a realistic and important set of channels and dynamics which factor into the value of medical breakthroughs.

A fourth strength of our macroeconomic approach is its ability to consider both the short- and long-run returns from advancements in aging science. We incorporate population dynamics and lags in therapeutic effectiveness. If an investment in medical R&D happens today, it is reasonable to assume that the benefits from this research would come years in the future. Furthermore, conservative rates of adoption across the population might suggest that those rates of effectiveness should be slowly phased in over further future years. We borrow from the literature on chronic medication adherence, assuming a 50% adoption rate, similar to that of existing chronic medication adherence (e.g. statins).^12^ Finally, demographic effects from changes in fertility and mortality rates can take decades to produce their full effect. Our macroeconomic model, with calibrated population dynamics, captures these and other changes in the economic environment, and the resulting values over the short and long run.

These features make our macroeconomic approach ideal to value benefits that only begin accruing decades in the future (as in the case of slight increases in fertility rates, enabled by slower reproductive aging). Our macroeconomic approach allows us to phase in the demographic and GDP outcomes of an advancement both annually and over several decades and to incorporate various assumptions about therapeutic adherence and adoption rates—all while the underlying population is realistically evolving according to true population dynamics. In our view, valuing improvements to a population’s aging profile requires a dynamic time series of the economy over the short and long run, incorporating the work, leisure, and productivity of the underlying population and businesses. This requires a macroeconomic model.

We use the open source OG-USA macroeconomic model of the United States in each of our simulations.^13^ This model supplements the existing literature in some important ways. VSL is the foundation of most estimates of the value of medical breakthroughs, and it is estimated in quasi-experimental settings, measuring individuals’ and governments’ willingness to pay for reductions in accidental death using labor market, housing market, and job injury data.^7,8^ The most common method in the literature involves calculating a QALY from either survey data or from a partial equilibrium life cycle model of an individual that is calibrated to have the total lifetime value of the researcher’s chosen value of a statistical life. The QALY approach from life cycle models uses a calculated multiplier on each year of life in the partial equilibrium model as the quality adjusted value of a life year, then shows how these values change with a medical breakthrough.

Murphy and Topel^9^ provide the seminal paper using a life cycle model calibrated with VSL estimates to estimate the economic benefits to longevity. They find that eradicating cancer would result in $84.7 trillion in economic benefits in 2025 dollars. Closest to our research question is Scott, et al.^10^ who use the approach of Murphy and Topel to estimate the value of increasing healthy life expectancy by one year and find a benefit of $46.6 trillion in 2025 dollars. Budish, et al.^11^ incorporate an average QALY estimate value into their model of strategic firm R&D behavior to estimate the net present value of reducing commercialization lags among pharmaceutical companies. Key assumptions in the approaches using the VSL or QALYs are the form of the household utility function (how individual welfare is modeled), the assumed VSL or QALY value, and the discount rate used to calculate net present values.

Two strengths of the life cycle model approach include (i) that it is derived from a well-defined theoretical value which directly captures individual welfare, and (ii) it is calibrated to match a VSL value derived from actual data and behavioral observation. However, the partial equilibrium approaches do not account for the heterogeneity across individuals in a population and the effect of those interactions across a long time series. When it comes to the specific research question of slowing biological aging, QALYs would also risk conflating “health” with “biological youth.” For instance, a 70-year-old suffering from just one disease of aging may be considered “healthy,” since most 70-year-olds suffer from at least two such diseases. Similarly, women in their 50s will predictably have lost the ability to give birth, and through a QALY lens, they may score a full point. This means QALYs can miss the fact that aging itself is a form of health decline, even absent clinical diagnoses.

Our modeling approach acknowledges that health and youth are separate processes, even if they strongly correlate. One can be 85 and “healthy,” or 25 and “unhealthy.” By looking at mortality, productivity, and fertility rates—and their stunningly predictable relationship with age—we bypass the problem of confounding health with biological youth. In other words, the world we get to paint with this model is not one where 70-year-olds are told they are healthy “for their age.” It’s a world where 70-year-olds suffer from the effects of aging observable in the economy at the rate of biologically younger adults. This includes, for instance, the ability to perform well in jobs with high cognitive demands. Our macroeconomic model assesses the effects of age-specific shifts in mortality, productivity, and fertility rates, capturing more subtle changes experienced by each age cohort. It also captures empirical rates of *voluntary* labor supply by adults ages 65 and older—an important factor when discussing the effects of slowing biological aging. Extensive online documentation and policy use cases exist for our open source model, the OG-USA overlapping generations macroeconomic model of US fiscal policy and demographic dynamics.^14^

Little research exists on the probability that investments will translate into breakthroughs.^15,16,17,18,9,11^ This is the case, not least, because each future breakthrough is unique and by definition has not happened, which means cost precedents are necessarily imperfect. To inform our simulations, we interviewed 102 scientists. Only 7.6% of the scientists who filled out our survey believe a 5-year delay in biological aging (five times as ambitious as the 1-year shift we simulated) would require more than $50 billion in funding, and 45.7% believe this could be proven with as little as $1 billion in funding, since funding *direction* is more important than funding *amount*. We simulate a maximum 1-year shift backwards in biological aging. The vast majority of scientists interviewed believe that any plausible investment amount required for these R&D advancements (in the billions) would be dwarfed by the returns on investment (in the trillions).

In early versions of this study, we assumed $50 billion in public investment for each R&D advancement; yet there were no material macroeconomic effects of the spending: the interest rate did not appreciate more than a few basis points, despite increases in government debt to finance this expenditure. Partly because of this, we do not include hypothetical investment costs in our simulations. Another reason why we omit specific investment costs is that higher investments in healthcare can mechanically increase GDP in the short run. This is a valid critique of GDP as a methodology, since it can manufacture the illusion that a given R&D area that is more difficult to engineer (requiring, say, $100 billion in funding) would result in higher returns in the short term than an R&D area that needs $1 billion. Because of these two points, and because R&D investments for medical research at the national level are usually in the hundreds of millions of dollars annually (while our macroeconomic results are in the hundreds of *billions*) we chose to ignore the costs of investment and probability of breakthrough discussion in this article.

## Results

We demonstrate our results from simulations in four key areas of aging research: brain aging, reproductive aging, and advancements in existing and potential near-future tools and therapeutics that may improve our ability to slow overall aging. In each case, we define how we map the particular medical breakthrough to the parameters of our model. In particular, we define specific correspondences between breakthroughs, shifts in mortality rates, fertility rates, and productivity levels by age.

In the first four simulations shown in Table 1, we assume a research-to-commercialization lag of 10 years, and another 10 years for maximum adoption. We consider a maximum adherence rate of 50% to reflect a foreseeable lack of universal access to novel therapeutics. The assumption of 50% follows the adherence rates to statins, a treatment that requires continuous use to ensure benefits.^12^ In Table 2, we consider the economic effects of achieving the endpoints of the TAME trial (Targeting Aging with Metformin) with any existing, FDA-approved therapeutic (e.g. GLP-1 agonists) and extrapolate on the future possibility of developing a therapeutic that achieves the same effects for still-healthy adults. For our first-generation therapeutic, we assume effectiveness begins in 5 years (2030), reaching full efficacy in another 5 years (2035).

**Table 1.** OG-USA simulation results for near-future advancements in aging science.

| Sim # | Simulation Description | Mortality decrease in years | Productivity increase in years | Fertility increase in years | Annual change in US GDP (Avg. over 2045-2064) | NPV of increased GDP over several decades | Increase in 2050 population |
| --- | --- | --- | --- | --- | --- | --- | --- |
| 1 | Slowing brain aging by 1 year | 0.20 | 0.70 | 0.0 | \$201B | \$8.9T | 268K |
| 2 | Slowing reproductive aging by 1 year | 0.02 | 0.10 | 1.0 | \$9B | \$9.3T | 391K |
| 3 | Slowing overall aging by 1 year for all adults age 40+ | 1.0 | 1.0 | 1.0 | \$408B | \$27.1T | 1.7M |
| 4 | Slowing overall aging by 1 year for all adults age 65+ | 1.0 | 1.0 | 1.0 | \$326B | \$14.3T | 1.2M |

**Table 2.**
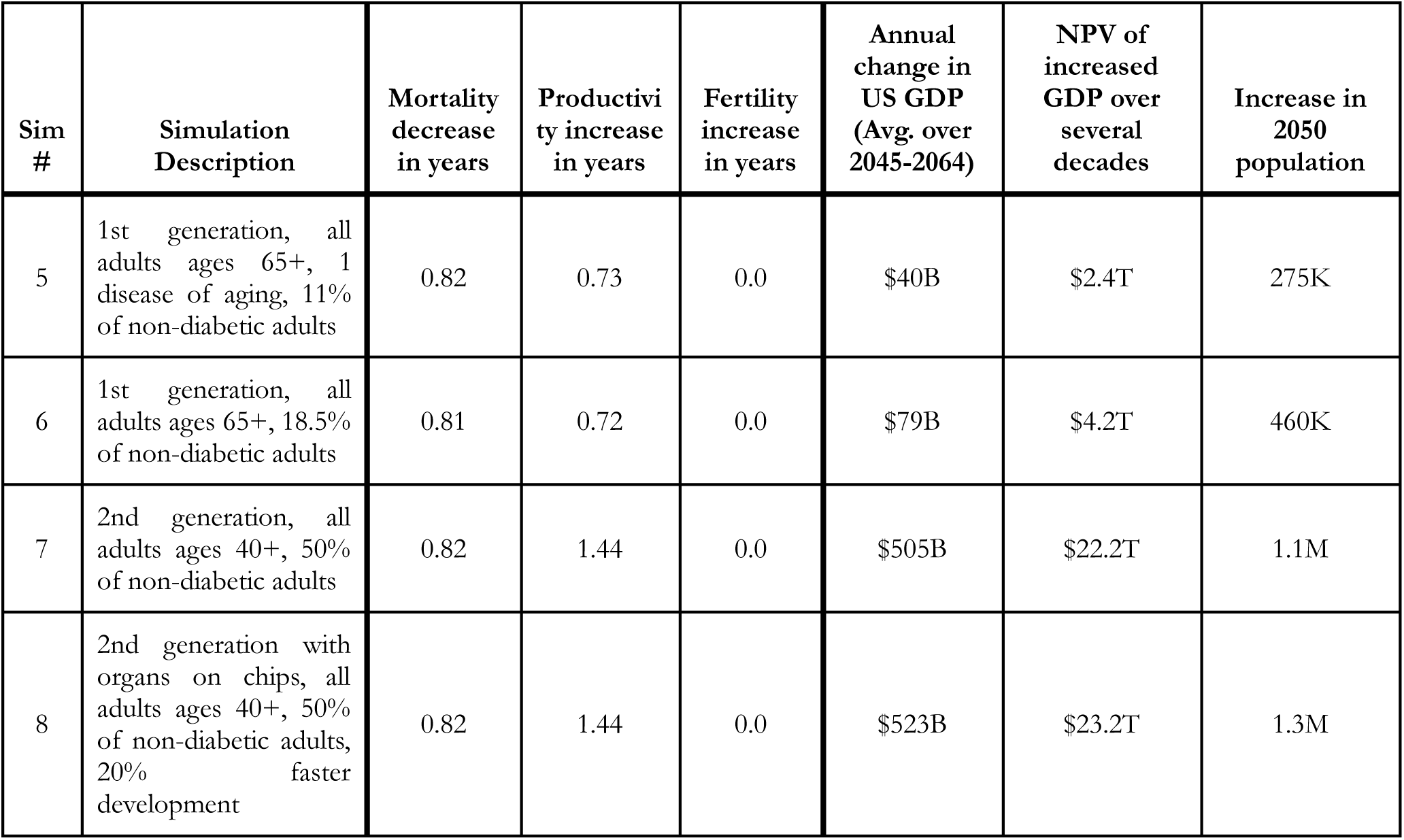
OG-USA simulation results for advancements in our ability to *measure* biological aging.

Table 1 lists our first of two sets of simulation results with their respective specifications. See Methods and Figure 5 for the results of sensitivity and scenario analyses.

### Slowing brain aging by one year

At its most compelling, aging science would enable full health extension, so that the function of all organs would be rejuvenated simultaneously. This, however, is unlikely to happen in the immediate future. And if one organ must be prioritized for rejuvenation, it is the brain. Past age forty, the human brain begins to shrink at a rate of up to 5% a decade.^19^ Processing speed, memory, and reasoning capabilities steadily decline. Though a distinction is often made between “healthy” brain aging and age-related neurodegenerative diseases, the boundary between normal and abnormal neurodegeneration is blurred. At age 65, “less than 5% of the population has a clinical diagnosis of Alzheimer’s disease, but this number increases to more than 40% beyond age 85.”^20^ Yet today, only 8% of the National Institute of Aging (NIA) budget is devoted to the biology of aging.^21^ To uncover new ways of intervening in brain aging, resources must be allocated to advance our understanding and measurement of the predictable and “normal” decline of cognitive function.

To simulate the effects of a 1-year delay in brain aging on the US economy and population, we assume these investments would pay off primarily by increasing the labor productivity of older workers and, to a lesser extent, decreasing mortality rates with age. We arrive at the year-change values for mortality and productivity rates in Simulation 1 from two datapoints. First, we refer to work by Lendqvist et al. to infer that roughly 70% of all labor productivity is brain-related.^22^ Second, we consider that 20% of all deaths can be attributed to the aging brain. Strokes and Alzheimer’s alone add up to 9% of all deaths in the United States, and their primary risk factor is aging.^23^ Parkinson’s, Amyotrophic Lateral Sclerosis (ALS) and brain tumors make up another 5% of all deaths. Some of these conditions (e.g. strokes) do happen in young patients occasionally, or in older adults as a result of causes unrelated to biological aging. The exact percentage of deaths attributable to the aging brain is impossible to estimate with today’s science. What is certain is that brain health is critical to overall health, including the proper regulation of metabolic and hormonal health, as well as the immune and nervous systems. Given the critical role of the brain in sustaining human life, we believe our assumption (that 20% of all deaths owe directly to the aging brain) is reasonable. Our baseline Simulation 1 considers a 0.2-year decrease in mortality rates and a 0.7-year increase in the age profile of productivity. We invite the interested reader to use our interactive simulation tool to use their own inputs (e.g. that only 60% of all labor productivity is brain-related.)

Considering the baseline assumptions above, we estimate the ability to extend the healthspan of the brain by just 1 year could add roughly $201 billion per year to US GDP, adding up to $8.9 trillion to the US economy in the long run (net present value), while extending 268,000 lives. Increases in labor productivity over a large population produce large aggregate effects. This, in turn, encourages labor force participation and capital accumulation. Inputting any plausible assumption on the role of the brain in human mortality and productivity, the socio-economic gains from this one-year shift in the cognitive healthspan of US adults aged over 40 would be significant. Healthier brains would mean more dignity to would-be neurodegenerative-disease patients and their families, and more productive, active years for older adults who *choose* to remain in the workforce.

Here, we also invite the reader to consider the contingency that perhaps due to prohibitive drug pricing, only 10% of the U.S. population over the age of 40 would benefit from a therapeutic that delays brain aging by 1 year. This low-uptake scenario would yield a still-significant $38.4 billion annual return to U.S. GDP, yielding $1.8 trillion in net present value, while extending 54,000 lives. For questions centering on access to novel therapeutics, timelines until market entry, therapeutic effectiveness, and even age cohorts who would benefit from a therapeutic, we encourage the reader to use our interactive simulation tool at silverlinings.bio, where readers can input their own (more or less optimistic) assumptions for each simulation.

### Slowing reproductive aging by one year

Evolution typically optimizes for reproductive health. From this, it follows that menopause is a relative rarity in the animal kingdom. Animals that experience no decline in fertility throughout their lives — like the naked-mole rat^24^ — are on one extreme end of the spectrum. *Homo Sapiens* are on the other extreme end. Most non-human primates continue to reproduce well into advanced age. In humans, however, the ovaries are the first organ to age — and men’s reproductive healthspan, too, declines more quickly than most assume.^25, 26^

Not all couples would choose to have children in their 40s if they could healthily do so. Yet at least 11% of females seek fertility treatments at some point in their lives, mostly due to age-related infertility.^27^ Importantly, this research area would do more than just increase birth rates. The health and social costs of ovarian aging are impossible to simulate fully in dollars, but even a narrow measure of its effects on GDP produce staggering results. In 2024, most women in the US choose to have children in their 30’s; yet given the current human healthspan, undergoing a geriatric pregnancy is both costly and dangerous.^28^ We do not yet understand what causes the female fertility window to last roughly 30 years less than the male window; why the reproductive span correlates with lifespan; or how aging ovaries influence overall aging. What is certain — given precedents in animal models like American lobsters — is that a short fertility window is far from a prerequisite for late-life health.^29^

Therapies which safely extend the reproductive healthspan would positively affect the health of older adults, and over several decades, they would increase the number of working-age adults. It is possible that the major gains from improving reproductive aging may come not from increased birth rates, but from the overall extension in healthspan accompanying delays in menopause.

Simulation 2 shows that a modest increase in fertility rates — on the order of about 1% — can produce significant economic effects in the long run. We model a one-year delay in reproductive aging as resulting in a 1-year shift in fertility rates by age, accompanied by a 0.02-year shift in mortality and a 0.10-year shift in productivity rates. The slight decrease in mortality and increase in productivity come from evidence such as that in Conti et al.^30^, who document the relationship between women’s reproductive health and labor productivity. In fact, Conti et al. note a more pronounced decrease in labor productivity rates by age after menopause, at roughly 1% per year within the first four years following an untreated diagnosis.) With these baseline assumptions, we find that slowing reproductive aging by one year would add $9 billion annually to US GDP in the short term, adding up to $9.3 trillion over several decades. The reader will notice that the long-run returns from this simulation are higher than the returns from slowing brain aging by one year, while the short-run returns are far smaller. Importantly, slower brain aging would afford near-immediate, vast returns to the economy. Slower reproductive aging would have some near-immediate impacts on increased productivity, but its most pronounced benefits in GDP terms would take place several decades in the future.

Slower reproductive aging leads to substantial increases in population: by 2050, our simulations show an increase of about 385,000 people. Roughly 26,000 of this population increase is due to lower mortality rates from conditions like menopause and the diseases downstream of it. The more than 350,000 new lives are not working-age adults, and therefore are not directly valued in the account of GDP. This demonstrates a potential weakness of GDP as a methodology, since it does not value the welfare gains of new lives for expecting parents and newborns themselves. Using a VSL methodology, newborn life years can be measured as immediately having economic value. Both methodologies are helpful. Our approach has been to supplement the existing literature by measuring more direct impacts on the US economy, in the short and long run.

From a GDP standpoint, more pronounced year shifts in reproductive aging would have diminishing returns. If reproductive aging could be reversed by 10 years, the fertility rate would not increase tenfold, even if the healthspan gains from slowing reproductive aging (e.g delaying menopause) would be significant. In other words, there are diminishing returns to how many more children women would have even if, like the American Lobster, female humans could safely reproduce through age 90. In the United States, again, about 11% of women seek infertility treatments throughout their lifespan.^27^ An even smaller percentage of couples who choose *not* to have children arrive at this decision due to age-related biological inability. Our simulation takes into account how the vast majority of U.S. couples would not have more children simply because, biologically, they could.

By contrast, therapeutics designed to slow overall or brain aging are less likely to suffer from this type of diminishing return. As we show in the appendix, slower reproductive aging would also lead to a *temporary* reduction in GDP per capita. This is partly because, as we introduce in the introduction of this paper, newborns take several decades to produce more than they consume. In the very long run, however, the returns from slowing reproductive aging are extraordinary, since they constitute at once investments in existing (e.g. would-be menopause patients) and future humans (e.g. otherwise unborn children to aging parents). Increased birth rates are unique in that new humans engender more new humans. Over several decades, newborns are likely to themselves produce more newborns, who then produce more newborns. Higher fertility rates have compounding effects we do not get by simply extending the cognitive healthspan of a living human.

All this means that to decide which simulation produces the most desirable returns (in terms of lives extended and GDP growth), one’s discount rate over the future is instrumental. While successful investments in the aging brain would offer near-immediate returns, over a multi-decade time series, investments in ovarian aging would offer slightly higher returns.

### Slowing overall aging by one year: making 41 the new 40, or 66 the new 65

At its most compelling, aging science would unlock not just functional gains for discrete organs or tissues, but organism-wide benefits. It is technically possible that a therapeutic might target the key causal node linked to aging across the human body. Yet advancements in organ, cell, and tissue transplants may be needed to unlock a holistic improvement in biological aging in the near future. For this simulation, we assume that cells, tissues, and organs in the body can be therapeutically targeted or replaced to result in a one-year delay in biological aging. This is different from extending life expectancy by one year. A one-year increase in life expectancy has been engineered many times before.^31^ Improvements in the biology of aging, even marginal, would have larger socio-economic benefits.

We simulate the effects of a convergence of therapeutics and technologies which delay biological aging — shifting mortality, productivity, and fertility rates — by one year for all US adults over age 40 (Simulation 3). Our model suggests that a mere one-year improvement in biological aging would increase US GDP by roughly $408 billion per year on average within the first 20 years of investment. If one considers the effects of this investment over several decades, the returns add up to $27.1 trillion in net present value to the US economy.

Because we assume no shift in retirement ages, a 1-year extension in healthy life would add slightly to Social Security costs. But adults aged 65 and older are the fastest growing labor group in the US, and this shift would substantially reduce medical costs while increasing labor supply. Our results in fact show that the increase in output (and thus income) generates more tax revenue than it does additional outlays due to increases in Social Security retirement payments. The returns also remain overwhelmingly positive for a 5-year shift backwards in biological aging using our economic model.

We assume the baseline group of adults affected is the population aged 40 and older. If, however, we limit our one-year shift in mortality rates, fertility rates, and worker productivity to the population aged 65 and over, the resulting economic value is smaller as it would consist of mostly *treating* older adults rather than *preventing* age-related decline in still-healthy ones. Yet even here, the results remain large (Simulation 4). This latter simulation shows an increase in US GDP of $326 billion per year within the first 20 years of efficacy, and a net present value of $14.3 trillion. Notably, all simulations offer substantial positive effects, even if all of their direct impacts on lifespan, productivity, and fertility are jointly reduced by 20%, as in our pessimistic scenario for each of the above. (See sensitivity analyses in Methods.)

### Slowing aging through existing tools, therapeutics, and new validation of biomarkers

Even if an existing therapeutic can improve aging with minimal side effects, proving this remains difficult. Because trials also typically measure gains in single diseases, the aging field suffers from the undervaluing of existing therapeutics that may improve biological aging. We consider the economic value of two drug classes. The first class is what we call “first-generation therapeutics.” These are existing, FDA-approved therapeutics that may marginally slow biological aging, but whose geroprotective effects have not been rigorously tested. The second class is what we call “second-generation therapeutics,” assuming that new drug targets can be found to safely delay aging in still-healthy, working-age adults. Put simply, the first simulation is a form of aging *treatment* for adults already suffering from at least one disease of aging, while the second is a form of aging *prevention*.

First-generation therapeutics are widely used as FDA-approved interventions (e.g. metformin, GLP-1 agonists, and rapamycin) with untested effects on the biology of aging. Today, they are most often prescribed to older, sick, or overweight adults *already* suffering from the diseases of aging. To simulate the economic value of administering a first-generation drug to slow biological aging, we consider the endpoints of the Targeting Aging with Metformin (TAME) trial — what may be the first-ever trial to measure geroprotective effects in humans with an imperfect, but well-studied drug.^32^ We assume the same (or superior) effects could be achieved by alternative drugs, but chose to model the effects of TAME due to the well-established safety profile of metformin *for older adults*, and due to the trial’s thoughtful endpoints design. For our mapping of the trial endpoints, see Methods.

This first-generation drug, if administered to 11% of the population 65 and older for biological aging (in line with expectations from the TAME trial), could increase US GDP by roughly $40 billion per year in the short term and $2.4 trillion over several decades (See Simulation 5 in Table 2). If the endpoints of the TAME trial are successfully achieved, the trial could pay for itself nearly 1000 times over within 1 year, with an investment cost of $50 million. This $40 billion annual increase in GDP would not be captured by any single pharmaceutical company, and this creates a pressing rationale for public and philanthropic investment. These gains come from more workers due to lower mortality rates, and from a more productive workforce. If we consider wider adoption of the same first-generation therapeutic (50% of all non-diabetic older adults), annual gains to US GDP would increase to $79 billion per year, adding up to $4.2 trillion over several decades (Simulation 6).

Adoption rates reflect different assumptions on what population groups would clinically benefit from or have access to novel therapeutics. Above, we simulate returns for the expected adoption of metformin for biological aging, considering that only 11% of the population of adults over the age of 65 with one pre-existing condition of aging would benefit from these therapeutics.. In Simulation 6, we also offer up a more optimistic adoption rate, at 18.5% of the population of all (non-diabetic) adults ages 65 and older.

The role of first-generation trials in collecting and validating biomarkers of aging and endpoints cannot be overstated. These trials will also be essential to legitimize the possibility of targeting biological aging. Yet in the longer run, the goal should not be to wait until working-age adults have become frail and sick, but to prevent the burdens of disease and social care. This is what we assume the counterfactual, second-generation drug class below could do.

Simulation 8 (2nd generation boosted by human-relevant methodologies like virtual cells or organs on chips) assumes a faster timeline until market entry, of 8 years (rather than 10) to initial efficacy and another 8 years to full efficacy.

For Simulation 7, we consider the effects of prevention rather than mere treatment in US adults aged 40 and above. Specifically, we assume that a second-generation therapeutic could be developed to benefit adults with no preexisting conditions, who will nonetheless predictably go on to suffer from one of the several diseases of aging. For simplicity, we assume the same endpoints of the TAME trial, but with a favorable risk-benefit profile for still-healthy, working-age adults. Our assumption is that the population-wide effect of this second-generation therapeutic is an average decrease in mortality rates of 3.1%, and an increase in labor productivity of 1.5%.

We find this preventative drug type could increase GDP by about $505 billion per year, yielding $22.2 trillion in net present value to the US economy over several decades. The change between first-generation and second-generation therapeutic (the latter designed for healthy, working-age adults), is substantial, at roughly $465 billion per year (Simulation 7 minus Simulation 5). Considering the effects of this shift from first-generation to second-generation therapeutics over several decades, this delta is worth an extra $19.8 trillion in net present value to the US economy. This demonstrates the value of truly preventative medicine, designed to extend healthy life rather than *treat* age-related conditions.

In the graph below, we demonstrate the effects of these two therapeutic types on the growth of the U.S. population. This counterfactual, second-generation therapeutic we simulate would result in more than an additional 800,000 adults alive by 2050 relative to the 1^st^-gen therapeutic, and 1.1 million additional adults relative to the baseline world without either development.

**Fig. 1.**
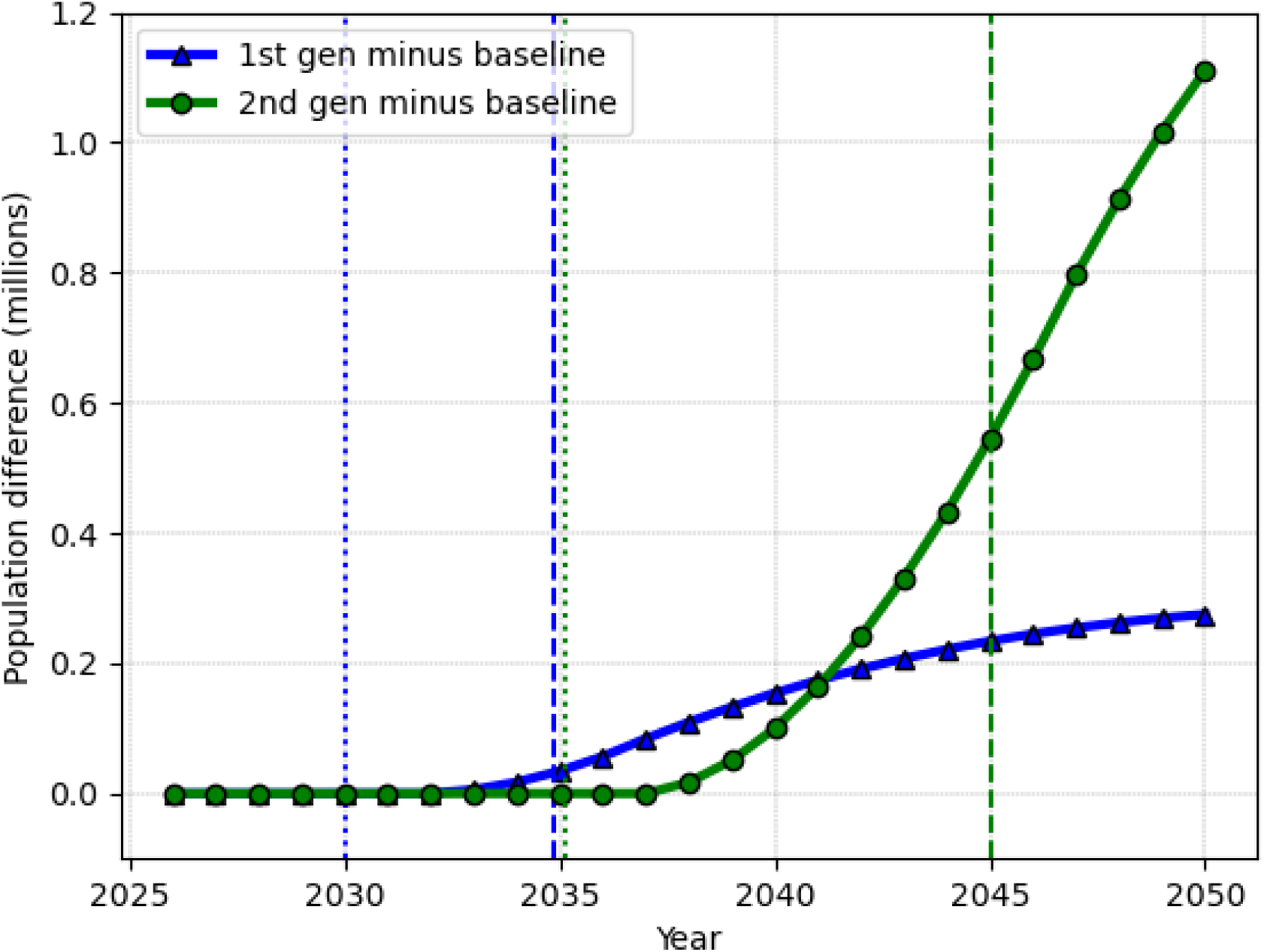
The effects of aging drugs on the U.S. population, for first- (existing, FDA-approved) versus second-generation (counterfactual, preventative) therapeutics. The 1st-generation therapeutic begins effectiveness 5 years after investment (2030), and reaches full effectiveness 5 years later. In the short term, existing, FDA-approved therapeutics extend more lives than our simulated 2nd-generation therapeutic. The latter simulation—of a counterfactual, preventative therapeutic—begins effectiveness in 10 years (2035), and reaches full effectiveness 20 years after investment (2045), saving more lives in the long run.

We acknowledge that arriving at a second-generation therapeutic would be a non-trivial achievement in drug development, moving away from treating diseases to slowing biological aging before clinical diagnoses. Developing a drug to delay biological aging with a favorable risk-benefit profile in still-healthy adults would be difficult. Yet most scientists we interviewed believe this is likely to happen in the coming decade. One recurring comment by scientists on what might enable this advancement was the ability to measure the effects of drugs more quickly *on humans*, instead of on other animals. This is especially important when it comes to biological aging, since no animal model ages exactly like humans do.

To simulate the economic effects of advancements in human-relevant methodologies like virtual cells (Simulation 8), which could accelerate translation, we consider the value of our second-generation simulation boosted by a 20% reduction in development timeline. This assumes a shorter timeline (8 years) to initial efficacy for our second-generation drug type. This acceleration in development alone could add another $18 billion to US GDP per year ($523 billion - $505 billion), or $1 trillion in net present value to the US economy in the long run ($23.2 trillion - 22.2 trillion). We consider this a conservative estimate of what human-relevant methodologies—ranging from new advancements in digital biology to validated organs-on-chips—could enable for aging research, since the same advancement could horizontally accelerate hundreds of clinical trials.

## Discussion

If the social ROI on aging research is so high, why aren’t more companies and research institutes investing in it now?

Well-documented biases in behavioral economics explain why people are often more willing to pay for a cure than for prevention.^33^ In turn, medical providers, pharmaceutical companies, and insurers have an incentive to overlook the value of health and longevity in favor of treating late-state or acute conditions.^34,35,36,37^ Insurers, hospitals, and patients face a tragedy of the commons whereby all parties would be better off prioritizing long-term health, but the incentives for individual agents to do so remain misaligned. For instance, because US citizens are free to change their health insurance at any time, few insurers invest in future patients’ health. And because governments largely subsidize our age-related health decline with programs like Medicare, we (and insurers) often underinvest in lifestyle choices like better diets — which can partly delay and even reverse *some* of the hallmarks of aging.

Government investment toward R&D targeting aging is also underprovided due to a generational misalignment. Because it can take years for R&D to translate into breakthroughs, the current population of older adults risks bearing all the up-front research costs without living to receive most benefits. This age group is more likely to vote and to exert a greater influence on politics. Today, it is often more profitable for a pharmaceutical company to lengthen the unhealthy life of a patient by a few months than to develop mechanisms that improve overall health. It is prohibitively expensive to test drugs that prevent dozens of diseases at once; it is easier to market “me too” drugs with little innovation than to develop new ones; and it is difficult to retain patents and patients in long clinical trials (which could quantify life-years saved).

Disease is often more easily measured than health. A therapeutic that preventatively extends the human healthspan is taken before its effects can be measured, and it compares against the unknowable counterfactual of how long the patient would have anyway lived in good health. All this leads to a system more prone to treating illness than preventing it, and to a backlog of undervalued therapeutics which, if tested and approved, could delay the onset of age-related health decline. The result is a lack of real preventative interventions, and an excess of drugs designed to treat the effects of aging at a point when they are hardly treatable.

Advancements which slow brain aging, extend the reproductive healthspan, or prevent the onset of multiple age-related, chronic conditions at once would have tremendous value to families and the global economy, with estimated returns in the trillions of dollars of economic activity and non-trivial population growth. Yet the current gap between health and life expectancy is unlikely to be closed by private markets alone, and this creates a pressing rationale for public and philanthropic investments.

## Methods

We summarize here three key components of our model, and how we integrate those into our simulations in this study: demographics, individual welfare and productivity, and government finances.

### Demographics

The OG-USA macroeconomic model has rich demographics. As shown in the following equations (1), the model simulates the behavior of a dynamic population of individuals of different ages across time 𝛚_s,t_ in which the number of individuals of each age is governed by a law of motion for the population based on fertility rates by age *f_s_*, mortality rates by age 𝛒*_s_*, and immigration rates by age *i_s_*. The time period is denoted by the subscript *t*, and the age group is denoted by the subscript *s*. Individuals are born at age *s*=1 and can live up to a maximum age *S*.

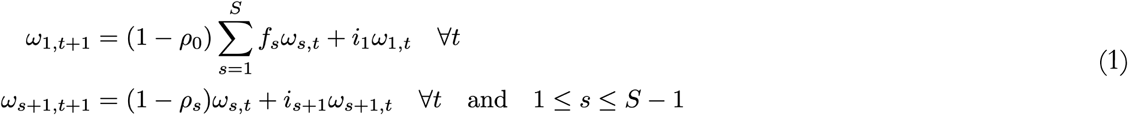

Mortality, fertility, and immigration rates are all assumed to be exogenous parameters of the model; they do not vary with income, wealth, or other endogenous variables. The mortality rates 𝛒*_s_* and fertility rates *f_s_* by age are key inputs to the model that we will change as being affected by aging therapy breakthroughs. Panels A and B of Figure 2 show US mortality rates by age and fertility rates by age. Figure 2 also shows our definition of a one-year shift in mortality and fertility rates. It is not a one-year shift in life expectancy, as in Scott et al.^10^ although our changes will certainly affect life expectancy. Instead, we are studying shifts in mortality and fertility rate schedules in which a one-year shift means that a 50-year-old now has the mortality rate of a 49-year-old, a 40-year-old has the mortality rate of a 39-year-old, and so forth. That is, we are directly modeling a one-year reversal in biological age. (Of note, 0 out of 102 scientists we interviewed responded that age reversal is impossible, even if there is no consensus on *how* it will be done.)

**Fig 2.**
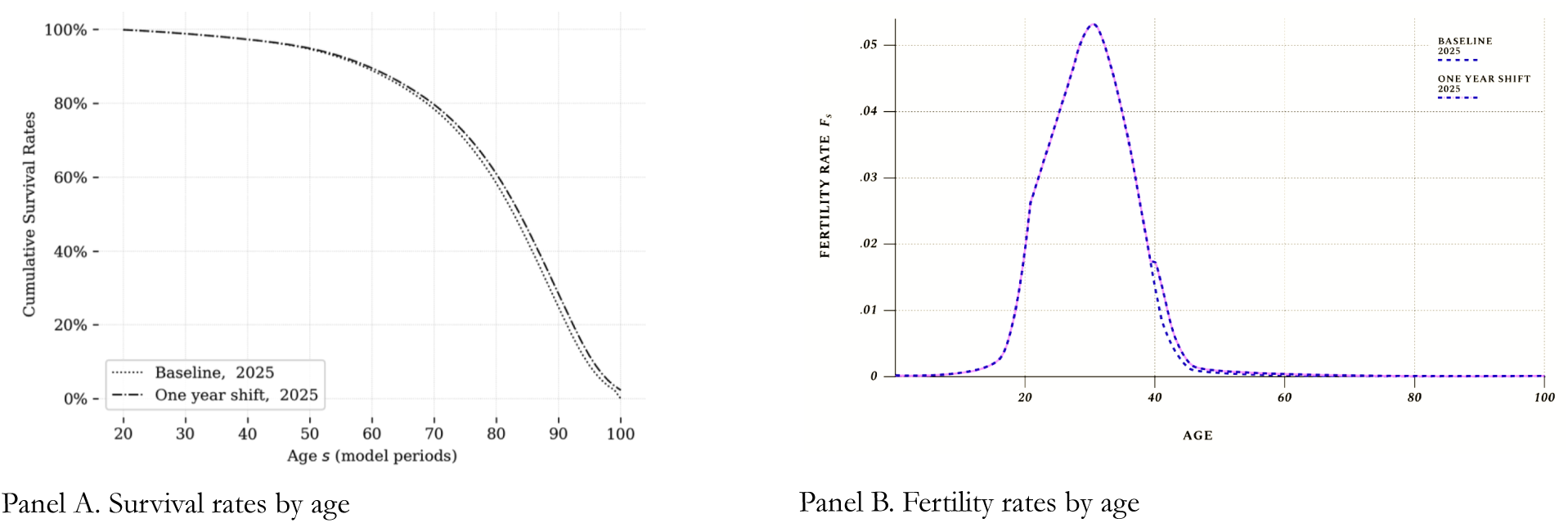
US survival rates and fertility rates by age, and a one-year shift for all individuals over age 40.

### Individual welfare and productivity

One way to think of the household side of the OG-USA overlapping generations macroeconomic model is as a group of individual life-cycle models in which new individuals are born every period, and a percentage of the individuals die at the end of each period. In each period a population of individuals are alive who differ in their age *s*, labor market productivity *e_j,s_*, and wealth *b_j,s,t_*. Each type of individual, with her different status, chooses how much to consume, save, and work every period and takes into account the future, her own survival probability, and what other agents might do.

Equation (2) is the per period budget constraint of an individual in the model, and equation (3) is a simplified version of the period utility function of the individual. On the left-hand-side of the equals sign of the budget constraint (2) are the costs of how a household might spend her resources: on total consumption *p_t_c_j,s,t_* or on savings for the next period *b_j,s+1,t+1_*. The right-hand-side of the equals sign of the budget constraint shows what are the resources or income of the individual. This includes capital income, labor income, bequests, remittances, government transfers, pension income and net taxes.

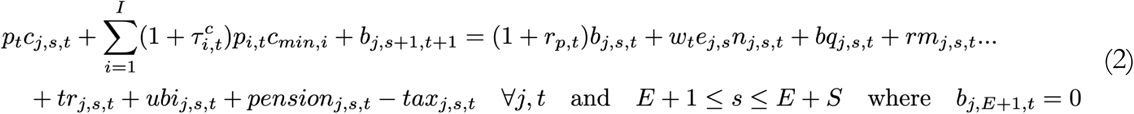

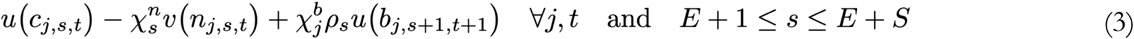

We focus on one part of the individual budget constraint (2) that is particularly salient to our simulations. The OG-USA model has carefully calibrated lifetime ability profiles *e_j,s_* that are part of the labor income term *w_t_e_j,s_n_j,s,t_* on the right-hand-side of the budget constraint in (2).These lifetime ability profiles are calibrated for 10 different lifetime income groups, which allows the OG-USA model to capture the income and wealth heterogeneity in the US. See “Lifetime Earnings Profiles” chapter of online OG-USA documentation for a detailed description of how the lifetime ability profiles are calibrated (https://pslmodels.github.io/OG-USA/content/calibration/earnings.html). We treat these individual specific labor productivity profiles as our variable for individual health. Thus when a medical breakthrough increases the productivity of an individual at a particular age *e_j,s,_*we increase those profile levels.

Figure 3 shows the calibrated lifetime earnings profiles by lifetime income group for the baseline in the OG-USA model and the effect of a one-year decrease in biological age individuals above age 40 such that the labor productivity of a 50-year-old becomes that of a 49-year-old, and so forth.

**Figure 3.**
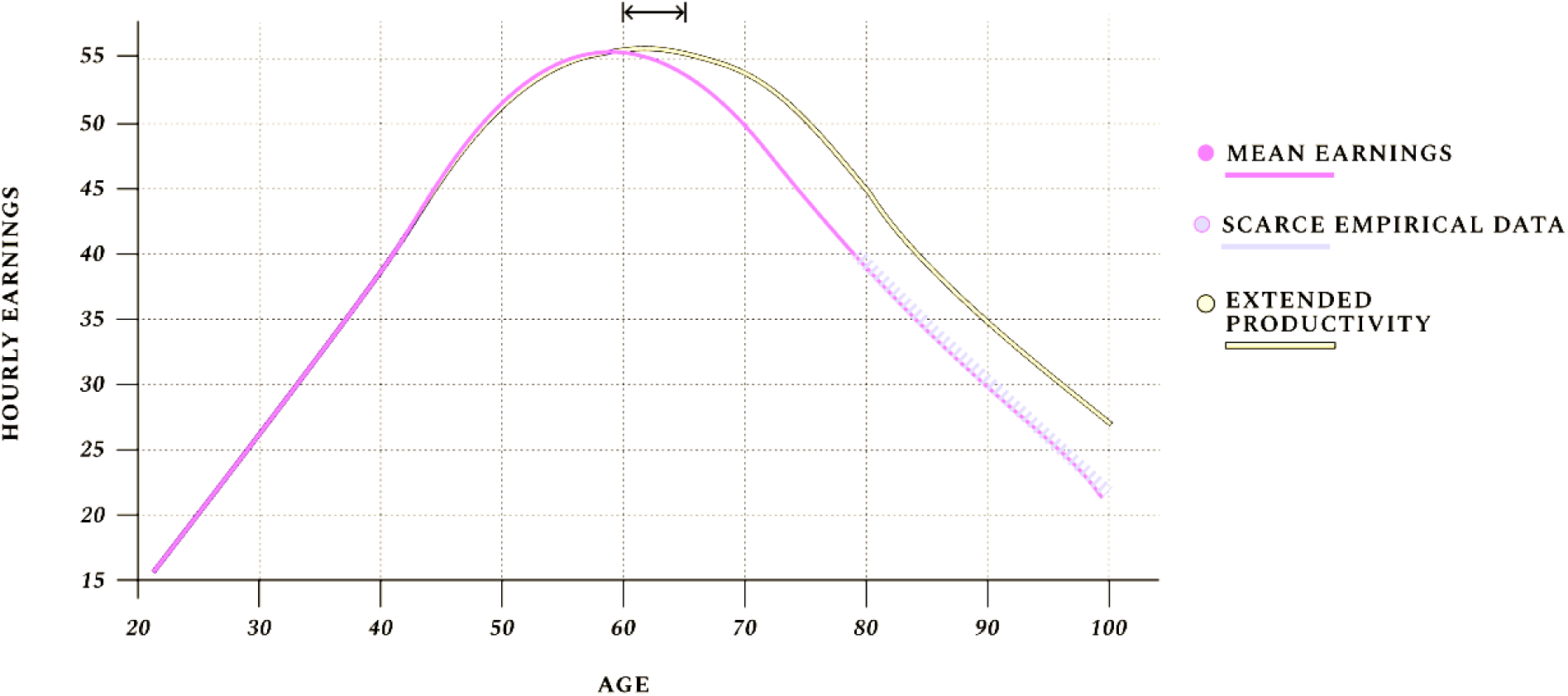
Lifecycle profiles of U.S. hourly earnings: baseline versus simulated 5-year shift in productivity rates by age.

Broadly speaking, people tend to reach peak earnings as long as their experience has not been outpaced by cognitive decline. This shows in the graph above, which documents how the average US worker records the highest hourly income she ever will around age 58, followed by a steep decline caused primarily by the effects of biological aging. In the average lifetime, hourly earnings increase as human capital goes up, then decreases with age-related health decline. In other words, human lifetime earnings follow a fairly smooth hump shape. We illustrate US mean earnings by age above, with a focus on hourly wages. Of course, older adults work fewer hours than younger ones, which means their total earnings are still often lower. Yet healthspan—and expected healthspan—is often the *primary* factor informing the number of hours people work. Changes in the biology of aging are unique because they can extend the average human’s productivity at the peak of this “triangle,” even considering only voluntary work after the age of 65. (Some readers may have noticed that the decline in hourly earnings between ages 60 and 80 look remarkably straight. It is a lucky artifact of the data that this decline *looks* linear. The ability profiles by age do *not* have a linear decline. Nor does the labor supply by age. Yet a lucky smoothing happens when we multiply hourly ability profiles with the average wage and labor supply, i.e. total hourly earnings).

The one other equation we highlight is the individual’s period utility function in equation (3). The individual’s utility or measure of welfare in every period is a function of how much she consumes (first term), how much she works (disutility of labor, second term), and a utility of bequests (bequest motive, third term). Mortality rates affect consumer decisions through their effects on the expected value of savings and bequests.

The term 𝝌^n^ multiplying the second term or disutility of labor of the period utility function in (3) vary by age and are calibrated so that the individual labor supply decisions for each age individual in the model baseline match the US labor supply data. As pointed out in Rupert and Zanella,^38^ this parameterization of the disutility from hours worked is critical to matching patterns of labor supply over the lifecycle. For example, without changing the disutility of work by age, older individuals in the model would supply much higher numbers of hours than observed in the data because hourly earnings do not fall very quickly, even after age 65 (see Figure 3). Whenever we increase productivity of individuals in the model *e_j,s_*as a result of medical breakthroughs, we also decrease the scale of the disutility of labor supply 𝝌^n^ . This is our definition of an increase in worker productivity. We find this simultaneous effect (changes in hourly productivity and in the disutility from work) an intuitive definition of worker productivity increases from medical breakthroughs. These assumptions imply that a slightly longer life lived in slightly better health would translate into slightly more hours worked. But we do not assume that hours worked grow linearly as biological age changes, rather hours worked are responses to the changes in lifetime resources and preferences as a result of healthier aging.

The combination of mortality rate changes 𝛒*_s_* and productivity changes (*e_j,s_*, 𝝌^n^ ) captures the two-dimensional decomposition of the effects of medical breakthroughs defined in Murphy and Topel^9^ and Scott et al.^10^ of changes in longevity across time and changes in welfare within each period.

### Production and technological growth

The supply side in the calibration used for our simulations consists of a representative firm which produces output using a constant returns to scale, Cobb-Douglas production function. Firms use capital and labor as inputs to production. There is an underlying growth rate in a labor-augmenting technology. This growth rate is calibrated to 2%, the mean rate of growth in US GDP per capita since WWII. Note that the underlying long-run rate of economic growth is exogenous, but during the transition from the economy of 2025 to the long-run steady state, the actual rate of economic growth will deviate 2%. Changes in demographics or health status do not affect this long-run rate of growth, but do affect labor productivity and will affect the growth rate of the economy over the transition to the steady-state.

### Government finance

The OG-USA model includes government taxes, government spending, deficits, and debt, as well as the effects of the incentives those policies have on individuals and businesses. The increased cost of Social Security pensions and the changes in other tax revenue streams and benefit program payouts from increased longevity in our simulations are some of the outputs from our simulations. We leave more in-depth fiscal analysis of the effects of longevity to future work.

You can see some of the tax and benefit variables on the right-hand-side of the equals sign in the individual budget constraint in equation (2). The following equation shows how government debt and deficits evolve as a function of tax revenue, spending, infrastructure investment, pension payouts, transfers, and interest payments. Detailed documentation for the theory behind all the components of government is available in the OG-Core online documentation chapter “Government” at https://pslmodels.github.io/OG-Core/content/theory/government.html.

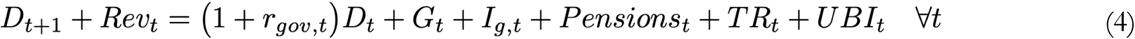

Pension amounts are determined through a direct modeling of the US Social Security system. Through our simulations, we hold the parameters of the Social Security system constant including the age of eligibility of old-age benefits, which we model as age 65. For this project, we deliberately simulated only marginal improvements in aging biology, with a focus on likely R&D advancements in aging science in the near future, based on interviews with 102 scientists. The most ambitious of our simulations models the impacts on the US population and economy of a 1-year shift backwards in biological aging. Our simulations show that increases in tax revenue through more productivity and marginal increases in work at older ages more than offset the increase in public pension outlays.

Even if governments choose to increase their debt or raise taxes to accommodate a growing number of retirees with the same benefits, reversing biological aging by one year would still result in lower medical costs, fewer unpaid caretakers, higher consumer spending rates, and extended careers for older adults who, in good health, would choose to remain in the workforce.

For readers interested in more substantial year shifts in biological aging, we recommend Goldman, et al,^39^ where the authors discuss how much the age of retirement may need to be raised to accommodate healthier aging. Yet we stress that in the near future, even just assuming voluntary labor supply of older adults, the more productive years enabled by investments in aging biology yield tax revenues that more than offset the costs of additional public assistance.

### Macroeconomic model, additional detail

To simulate the counterfactual futures discussed in this study, we begin by mapping changes in healthspan and lifespan from potential breakthroughs into changes in mortality, productivity, and fertility rates. These three levers directly influence people’s decisions and ability to work, save, consume, and live. The resulting behaviors, in turn, shape aggregate output, employment, and other macroeconomic outcomes. We focus on US Gross Domestic Product (GDP), which is directly impacted by the number of working-age adults and their productivity. We deviate from most approaches in the literature on the economic value of health and longevity and use what is called a "general equilibrium" model. This means we take into account the effects of supply and demand across people and markets (e.g., labor or assets) over several decades, which allows us to account for how aggregate economic outcomes affect individual decisions. For instance, people who know they have longer to live tend to invest more in their education, work more, and save more.

The model parameters are calibrated so that the model output matches a number of characteristics of the US economy. These include the capital-output ratio, the debt-to-GDP ratio, the distributions of wealth and income, variations in labor productivity across the lifecycle and across skill groups, the profile of hours worked by age, average and effective marginal tax rates on households and businesses, and other salient features. DeBacker et al.^40,41^ provide more details on the calibration.

#### Demographic Change

The model we use includes overlapping generations of US households who make decisions on consumption, savings, and labor supply over their lifetime. Additional details on the structure of the model can be found in the online documentation for OG-Core at http://pslmodels.github.io/OG-Core. The size of these cohorts of people and their chances of growing sick and dying before new technologies are introduced are all drawn from United Nations World Population Prospects (WPP) data on projected fertility, mortality, and immigration rates for the US. These households vary in their productivity levels, which also evolve as they age.

Figure 4 shows how the percent of the population at each age changes across time for a given initial population 𝛚*_0,t_*, mortality rates 𝛒*_s_*, fertility rates *f_s_*, and immigration rates is. It is one of the critical strengths of the OG-USA model that we can get these population dynamics right that result from changes in breakthroughs in aging therapies because the changes in population dynamics have large macroeconomic implications.

**Figure 4.**
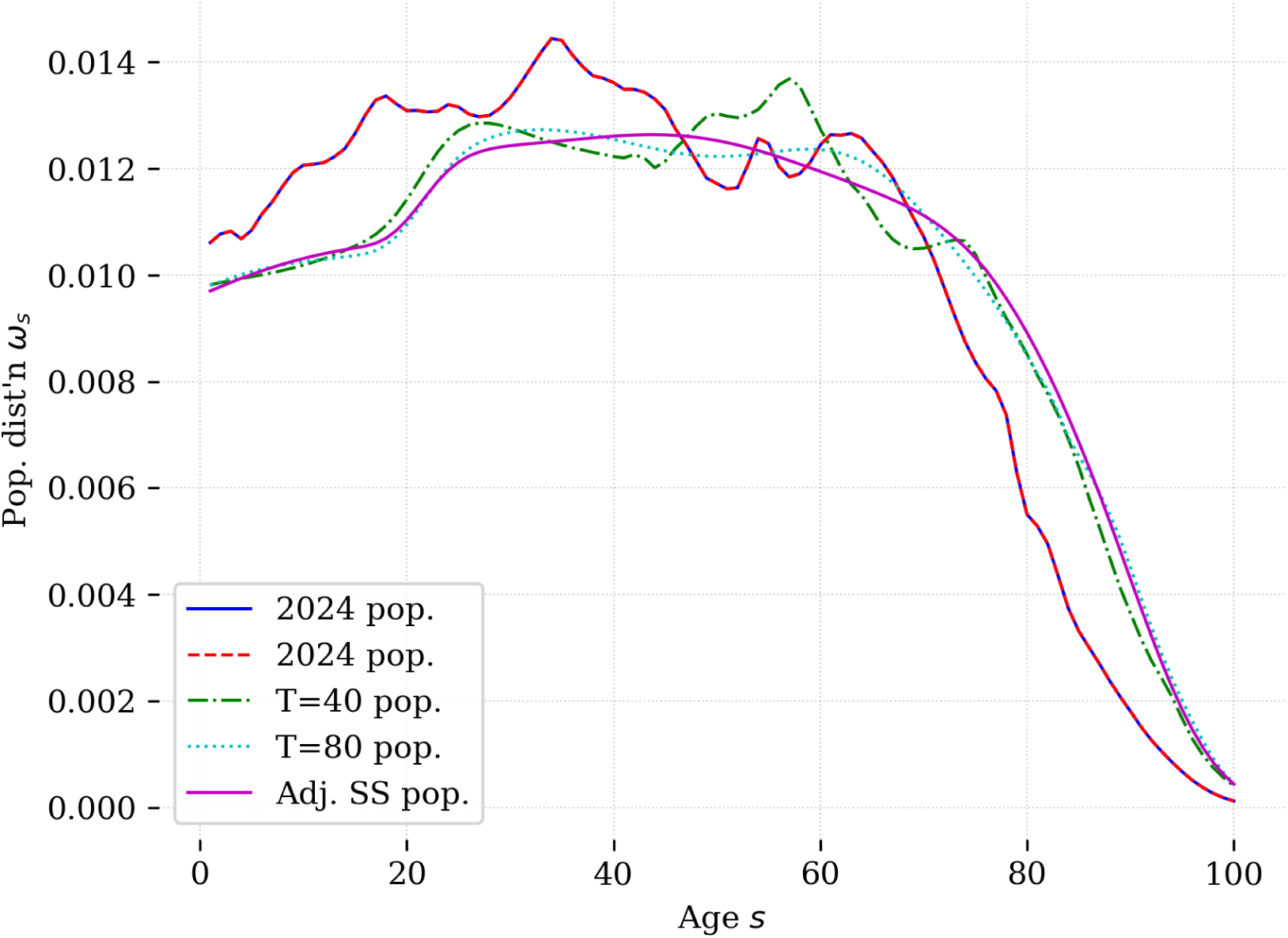
Evolution of the US population distribution over time: 2024-2104.

#### Biological Aging

In our life-cycle model, age serves as a key variable, and individuals’ decisions naturally shift over time as they grow older due to evolving labor productivity, accumulating savings, and a shortening of expected remaining lifespan. We incorporate the potential impacts of improved biological aging through three primary channels: (1) labor productivity and the corresponding disutility of labor supply, (2) mortality rates, and (3) fertility rates. By addressing aging, we assume that cognitive and physical decline is slowed, thereby enhancing *voluntary* productivity in later life and reducing both the direct and opportunity costs of working. Consequently, this lowers the disutility associated with labor supply. Furthermore, we posit that treating aging directly influences mortality by mitigating age-related disease and deterioration. Finally, we assume that treatments targeting biological aging will extend reproductive healthspan, leading to a modest rise in fertility rates, taking into account modern reproductive decision-making.

#### Productivity and mortality rates

In each period of their life cycle, households determine how much to save, consume, and work to optimize the discounted expected utility over their entire lifespan, subject to a budget constraint. The utility function for each period is represented as:

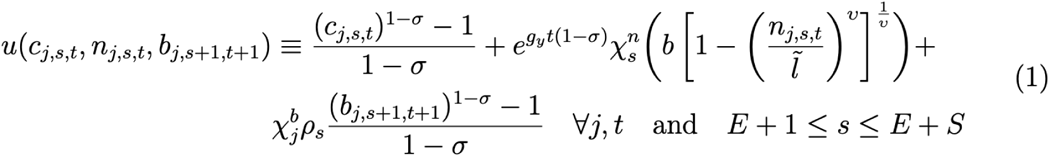

Consumption of the household of type j at age s in time period t is given as c_i,s,t_, labor supply is denoted n_j,s,t_, and household financial assets at the start of period t are b_j,s,t_, while their savings they leave period t with is b_j,s+1,t+1_. Note that consumption is strictly positive, while labor supply must be in the set (0, ̃l], where ̃l is the maximum fraction of a model period a household can work. The first component of the utility function is a constant relative risk aversion function of consumption, with a coefficient of relative risk aversion (CRRA) given by σ. The second component of the utility function represents the disutility of labor supply. This component has a utility weight of χ_ns_ , which varies by age s, representing changes over the lifecycle in the direct and opportunities costs of participating in market work. There is also a term in this component of the household utility, which is e_g_y_t(1−σ)_. This term is included so that the disutility of labor is scaled by the underlying economic growth, given by g_y_. This scaling is necessary because household consumption and savings grow at the rate g_y_, while labor supply is stationary. Thus, without this scaling, in the long run, the disutility of labor would shrink relative to the contributions from consumption and the bequest motive. The disutility function is an elliptical function, developed by Evans and Phillips^42^ It has the nice property of asymptotes at 0 and 1, which in turn helps the numerical optimization stay within the bounds on labor supply. The third and final term in the utility function is warm glow bequest motive. This term has a utility weight of χ_bj_, which varies by household type (skill level) in order to match the skewed distribution of bequests. Note that this term is discounted by the mortality rate, ρ_s_, since a household only leaves a bequest if they do not survive to the next period. The function for the warm glow bequest motive is also a CRRA function with the same coefficient as the function for consumption. The functional form for the consumption and bequest motive both share the same shape. This helps ensure the model can be stationarized when there is underlying technological and population growth that increases the size of c and b over time.

The household maximizes the discounted expected value of the lifetime of utility flows:

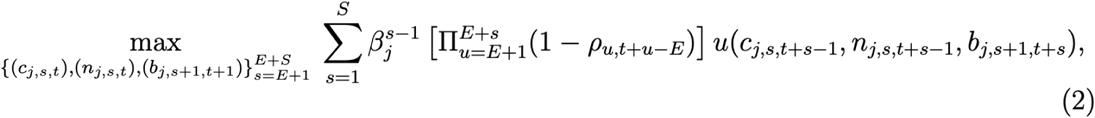

Where β_j_ is the discount factor for households of type j. Utility is maximized subject to the household’s budget constraint, where the period budget constraint is given as:

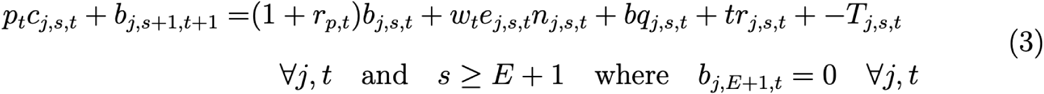

The budget constraint has the following prices. The price p_t_ represents the tax inclusive price of the composite consumption good in period t, r_p,t_ is the rate of return on the household’s financial portfolio, and w_t_ is the wage rate per effective unit of labor. The parameter e_j,s,t_ represents the labor productivity of households of type j of age s in time t and can be thought of as the number of effective units of labor supplied per unit of time working. Reading from left to right, the budget constraint says that in each period spending on consumption (p_t_ c_j,s,t_ plus new asset purchases (b_j_ , s + 1, t + 1) must equal gross asset income ((1 + rp, t)b_j,s,t_) plus labor income (w_t_e_j,s,t_n_j,s,t_ plus bequests (bq_j,s,t_) and government transfers (tr_j,s,t_), less total income and wealth taxes paid (T_j,s,t_).

One can use the budget constraint to solve for c_j,s,t_ as a function of saving (b_j,s+1,t+1_) and labor supply (n_j,s,t_), reducing the household’s per-period problem to the choice of savings and labor supply. The necessary conditions for the optimal choices of savings and labor supply in each period are given by the following equations:

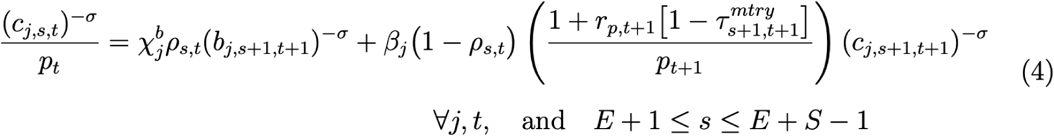

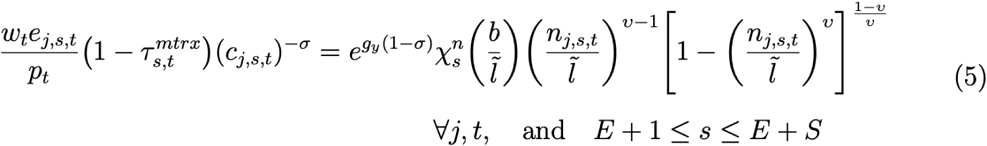

Equation 4 tells us that the household will choose savings to equate the marginal utility per dollar spent today to the expected marginal value of a larger bequest plus the discounted, expected marginal utility of consumption in the next period, where expectations are over the probability of survival, (1 − ρ_s_). Equation 5 notes that at the optimal choice of labor supply, the household will equate the marginal utility of consumption per hour worked to the marginal disutility of labor supply.

Within the household’s optimization framework, both mortality and productivity explicitly factor into the relevant equations, so any enhancements to aging will affect these terms. Focusing on mortality, it appears in the first-order conditions for savings. When mortality declines, households place a higher value on future consumption, boosting their savings. At the same time, however, the expected value of a warm-glow bequest diminishes, which dampens the incentive to save. Consequently, the net effect of changing mortality rates on savings—and thus the overall capital stock—remains ambiguous.

When it comes to productivity, we consider the lifecycle profiles of effective labor given by e_j,s,t_ as well as the utility weight on the disutility of labor supply, χ_ns_ . These show up in the household’s first order condition for the supply of labor, Equation 5. An increase in e_j,s,t_ will have two countervailing effects on labor supply. First, there is a substitution effect. Namely, if e_j,s,t_ increases, the household gets more income for each hour worked and will substitute away from leisure and towards market work. On the other hand, as e_j,s,t_ increases, household income goes up and the household will choose to consume more leisure. Thus, changes in labor supply from a change in effective labor per hour worked have an ambiguous effect on labor supply and will depend on the relative size of the substitution versus income effects. On the other hand, a change to the utility weight χ_ns_ affects labor supply in a clear direction. Lowering the disutility of labor supply, through a lower value of χ_ns_ , will significantly increase labor supply.

#### Mortality and fertility’s effects on the economy

We now examine how mortality influences the broader economy. Here, too, is where shifts in fertility have a visible impact. To illustrate these effects, we will briefly discuss how demographics are incorporated into the macroeconomic framework and the market-clearing conditions required for general equilibrium.

Ω_t_ represents the population distribution, the number of agents of each age s, at time t. This Ω_t_ matrix has elements ω_s,t_ and these ω_s,t_ evolve according to the law of motion:

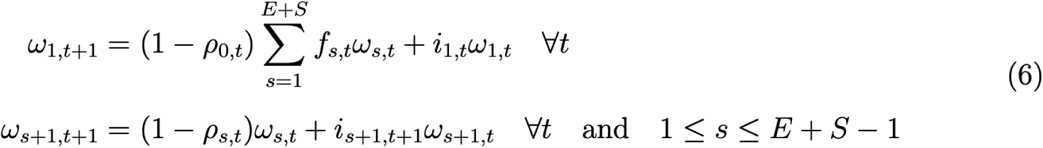

The terms in these equations not described above are f_s,t_, the fertility rates of agents of age s at time t, and i_s,t_, the rate of immigration of agents of age s at time t. Fertility rates are given as the fraction of agents of age s that have a child. Immigration rates are defined as net immigration of agents aged s as a fraction of the population aged s. The first equation in 6 says the number of one-year-olds is equal to the number of newborns (f_s,t_ω_s,t_) who survive infancy plus the number of net immigrants of age one (i_1,t_ω_1,t_. The second equation says the population of those older than one evolves as a function of the survival rates of those in the country plus next immigration. From these two equations we see that improvements in reproductive aging which result in increased fertility rates in turn increase the size and growth rate of the population. By contrast, decreases in mortality will increase the size of the population, but not its long-run growth rate.

In terms of the macroeconomic effects of mortality and fertility, it is the changes in the size of the population that will impact economic aggregates. The market clearing conditions perhaps show this most clearly. The market clearing condition for the labor market says that total labor demand from producers, L_t_ must be equal to the total supply of effective labor units from households:

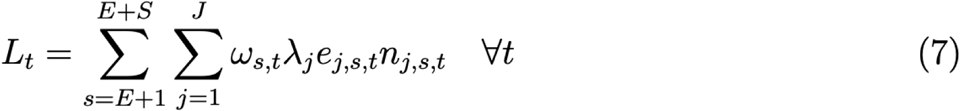

If the population grows—due to reduced mortality or increased fertility—then the total labor supply rises. In equilibrium, wages adjust to balance this expanded labor supply with firms’ demand. As more labor is employed, overall production increases according to the aggregate production function Y = F (K, L). Note also from this market clearing condition that holding constant labor supply, an increase in productivity (e_j,s,t_) will increase the aggregate effective labor supply and increase output. Similarly, for asset markets, the total demand for debt and private capital must be equal to the supply of savings from the household:

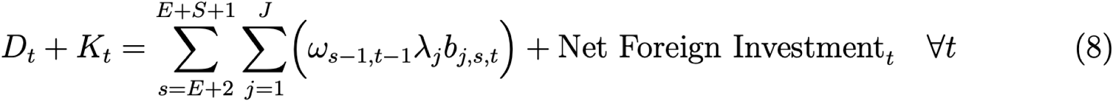

As well, lower mortality and higher fertility lead to a larger population and thus a larger supply of savings. This increase in the supply of savings will lower interest rates until the capital demanded (K_t_) and government bonds (D_t_) equal the supply of savings. More capital demanded by producers, in turn, increases output.

#### Quantifying the Effects of Slower Aging

We have identified the quantitative measures by which the three key levers—productivity, mortality, and fertility—are influenced in our simulations. Here, we explain briefly how these measures are implemented in our model.

When modifying any of these levers, we must make assumptions about the average size of the effect and how that average manifests across different age groups. In other words, we must specify how treatment impacts vary with age. At present, existing clinical studies provide limited benchmarks for average treatment effects targeting aging, mainly due to the market failures highlighted earlier, which have prevented trials from fully capturing age-specific outcomes. Although animal studies have produced notable findings, human trials on aging are still in their early stages and will require several years to complete.

Given this uncertainty, we approximate age-dependent effects by adjusting known age trajectories (for instance, the way mortality generally rises with age). More concretely, when simulating a change in one of the levers, we shift the observed age profile forward (or backward) so that it matches a particular average treatment effect. We thus assume that, if 41-year-olds were to have the same biological characteristics as 40-year-olds, they would face mortality rates consistent with those of current 40-year-olds, be as productive on the job as current 40-year-olds, and have fertility rates similar to current 40-year-olds. This approach is viable because we already possess robust data on how labor, consumption, and saving vary by age among the existing adult population.

Panel a of Figure 2, shown above, illustrates a one-year shift in mortality rates for individuals aged 40 and above, as reflected in the survival functions before and after our simulated shift. This technique helps approximate average treatment effects and aligns conceptually with aging research. By “moving” older individuals’ profiles so that they exhibit the mortality, productivity, and fertility patterns of people one year younger, we can infer how US adults would make decisions if their biological age were similarly shifted.

Similarly, we can think of shifting fertility rates to approximate a change in biological age. Panel b of Figure 2 shows US fertility rates in 2025 from the WPP data and a counterfactual one-year shift in fertility to represent one the the effects of slowing reproductive aging.

### Model Solution and Results

The model solution is made up of a long-run steady-state equilibrium, as well as a transition path equilibrium, which represents the economy from its present state in 2025 up to the long-run equilibrium. Solving for the economy’s full equilibrium path — namely, ensuring that the decisions of consumers and producers are consistent with one another in each year of the model’s simulation — allows us to analyze transitory effects of economic interventions, something that has been important for the simulations that followed.

For each model simulation, we provide the results summarizing the annual gains to US GDP as well as the net present values (NPVs) of those changes over a longer time horizon. Non-economists can think of NPVs as the long-term returns over several decades on present-day investments, understood through the value of a dollar in today’s economy. In every simulation, we estimate an annual dollar value of the estimated changes to US GDP, and calculate the NPV assuming a constant discount rate of 2%, consistent with the Congressional Budget Office’s long-term forecast.^43^

#### Sensitivity and Scenario Analyses

In this section, we provide sensitivity analyses of our results from Tables 1 and 2 as a way of exploring potential sources of uncertainty in our estimates. One of the most critical model parameters for the net present value calculations of the economic returns to slowing aging is the discount rate used. In these two tables in the paper, the assumed discount rate for our net present value calculations was 2%. In Table 3 below, we show the net present value calculations for varying discount factors 1%, 2%, 3%, and 4%. The net present value of the returns shrinks as the discount rate increases. More subtle is the fact that different simulations show different relative effects of changing the discount rate. Consider the comparison of slowing brain aging by one year and slowing reproductive aging by one year. At the baseline 2% discount rate, the NPV of the economic effects of slowing brain aging exceeded those of slowing reproductive aging. However, at a lower discount rate of 1%, the relative size flips; reproductive aging shows larger economic effects. This highlights how the discount rate interacts with the timing of the benefits of reduced biological age. Since the benefits of reproductive aging on GDP mostly come from a larger working age population as result of higher fertility, these benefits are decades in the future. Whereas the benefits from slowing brain aging are largely productivity gains which begin as soon as this intervention is available. In general, simulations where benefits are materialized further in the future are affected more heavily by using a higher discount rate.

**Table 3.** Sensitivity of the NPV of increased GDP to the discount rate.

| Sim # | Simulation Description | NPV of increased GDP over several decades |  |  |  |
| --- | --- | --- | --- | --- | --- |
|  |  | Various discount rates |  |  |  |
|  |  | 1% | 2%<br>(Tables 1 and 2) | 3% | 4% |
| 1 | Slowing brain aging by 1 year | \$16.1T | \$8.9T | \$5.2T | \$4.1T |
| 2 | Slowing reproductive aging by 1 year | \$20.7T | \$9.3T | \$4.3T | \$2.9T |
| 3 | Slowing overall aging by 1 year for all adults age 40+ | \$53.3T | \$27.1T | \$14.6T | \$10.9T |
| 4 | Slowing overall aging by 1 year for all adults age 65+ | \$26.0T | \$14.3T | \$8.4T | \$6.5T |
| 5 | 1st gen, all individuals age 65+, 1 disease of aging, 11% of non-diabetic adults | \$4.3T | \$2.4T | \$1.5T | \$1.2T |

Our model depends on a number of assumptions around people’s behavioral responses to therapeutics that extend healthy lifespan. As these therapeutics are mostly still under development (or have not begun development, as in the case of our second-generation therapeutic), our baseline scenarios are by necessity speculative. We therefore cannot fully characterize the distribution of uncertainty around each variable in our models, meaning that conventional Monte Carlo-based methods of uncertainty quantification were closed off to us. As a way to bound our estimates, we instead simulated optimistic and pessimistic scenarios for each of the major interventions we discussed. For each intervention, we produced a pessimistic scenario in which the impacts of all targeted outcomes (i.e., mortality, productivity, and/or fertility) were jointly reduced by 20% compared to their baseline value. In the optimistic scenario, we instead increased all impacts by 20% for each targeted outcome.

For simulations based on the TAME trial, we instead focused on the mortality impact alone since target ranges for this measure have been published. Specifically, we varied the hazard rate of mortality from 0.88 to 0.99 around a base estimate of 0.93.

Our scenario analyses show that even pessimistic assumptions on the impacts of these R&D advances still allow for these interventions to deliver substantial economic benefits. (See Figure 5.) There is more uncertainty, though, in the relative value of different interventions across different types of benefit. For instance, all scenarios show that improving brain aging offers more average GDP improvements than improving ovarian aging. However, there is overlap in how much benefit the two interventions offer in terms of net present value and population change. Our scenario analysis of the GDP impacts of the “66 is the new 65” scenario also overlaps in some scenarios with the estimated impacts of improving brain aging. We conclude from this that the “41 is the new 40” intervention is highly likely to offer the highest net present value of GDP change. We are less certain about which of the other three interventions is second-best. However, we can be reasonably certain that the “41 is the new 40” and “66 is the new 65” interventions increase population more than the brain and ovarian aging interventions. Our interviews further corroborate how these returns would dwarf any plausible investment amount.

**Figure 5.**
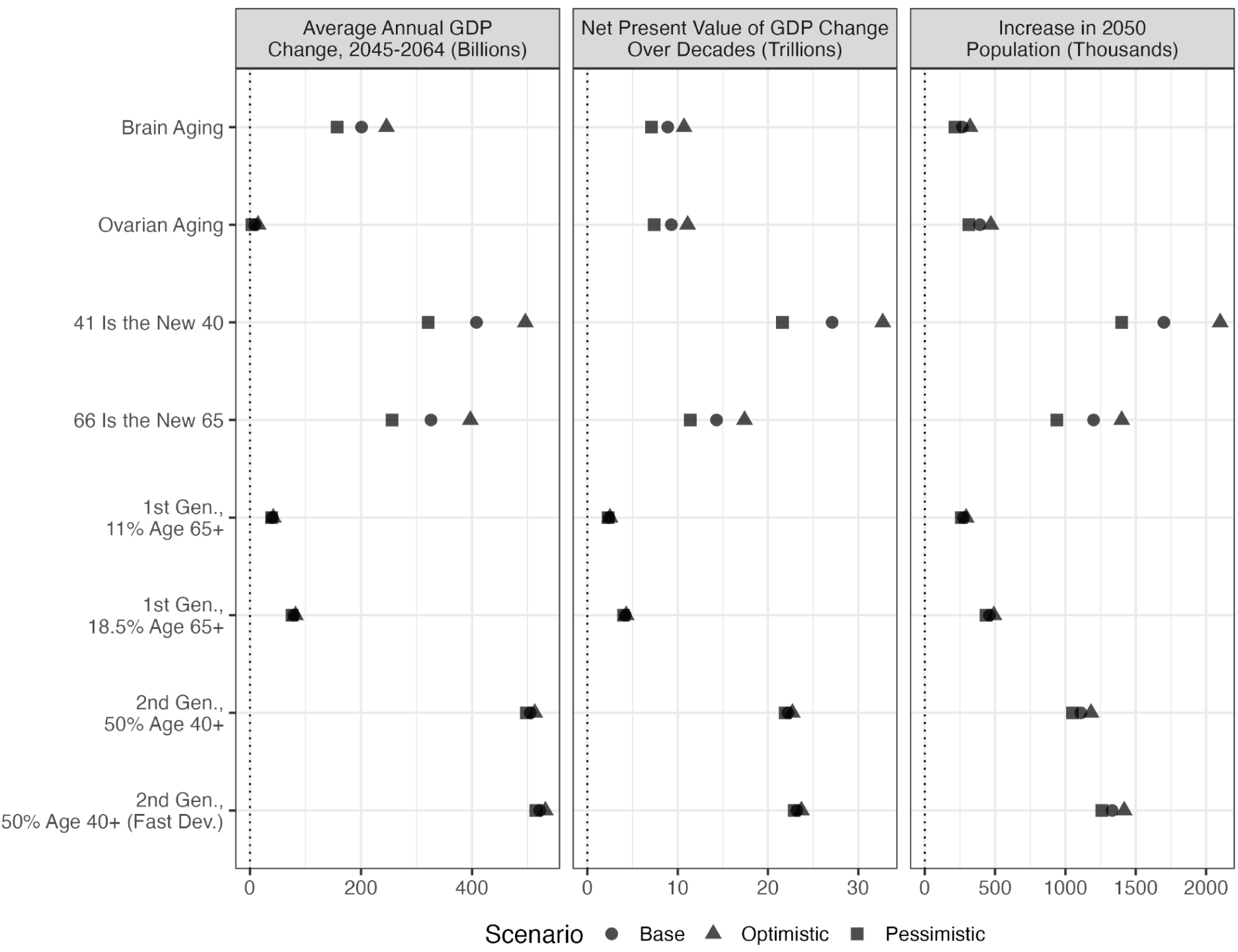
Estimated impacts of different interventions on GDP and population, with alternative results of scenario analyses.

### Measuring returns in GDP per capita

To illustrate the returns to GDP of each simulation on a per person basis, the table below summarizes the aggregate returns (in terms of a net present value over several decades) alongside the changes in GDP per capita over the medium term (2045-2064).

**Table 4.** Slowing Aging By 1 Year, GDP Per Capita Results.

|  | NPV<br>(effects on GDP over 100<br>years) | GDP PER CAPITA |
| --- | --- | --- |
| Slow brain aging by<br>1 year | \$8.9T | \$508 |
| Slow reproductive<br>aging by 1 year | \$9.3T | -\$5 |
| First-generation<br>therapeutics<br>(Existing,<br>FDA-approved<br>drugs repurposed<br>for biological aging) | \$2.4T | -\$20 |
| Slow overall<br>biological aging by 1<br>year | \$27.1T | \$425 |

The reader may notice that some advancements which are obviously good — like extending women’s reproductive window — have negative effects on GDP per capita in the short term even as they increase many people’s wellbeing and GDP. Why? First, because from a GDP standpoint, the newborns enabled by lower age-related infertility, miscarriages, or delayed menopause take over two decades to produce more than they consume. (As we note in the introduction, newborns don’t work, and they temporarily remove their parents from the workforce.) Second, negative GDP per capita results also occur when simulating near-future advancements premised on *existing*, FDA-approved therapeutics, more aligned with today’s sick-care treatments. We explain both instances below.

GDP per capita is negative for two of our simulations: slowing reproductive aging and what we call “first-generation therapeutics,” i.e. running a clinical trial similar to TAME, with an existing, FDA-approved therapeutic. In these two simulations, we choose to model near-future, narrow advancements in aging science that do not achieve the more optimistic and desirable goal of *preventing* age-related decline in still-healthy adults. TAME treats adults aged 65 and older with at least one preexisting disease (note our description of its sick-care nature, in contrast to our "2nd-generation" therapeutic), and slower ovarian aging produces more newborns with only small effects on the labor productivity of would-be menopause patients.

Of course, the ideal scenario for advancements in aging biology is to allow us to holistically target aging — such that *all* our organs, cells and tissues become biologically younger at once. This very optimistic scenario would produce strictly positive GDP per capita results, and we model it in the remainder of our simulations. But we trust this is likely *not* the scenario aging science will produce in the very near future. In the absence of extraordinary results in, say, partial reprogramming or sweeping gene or cell therapies that transform how we age as a species, advancements that improve the aging profile of a single organ (like the ovaries) or that *treat* older adults in imperfect health (like TAME) are very welcome — even if they *temporarily* decrease GDP per capita. This is the case for many of today’s drugs (e.g. cancer therapeutics that keep 75-year-olds alive in poor health), and we believe it is important to model these less optimistic outcomes of progress in aging science, too. Advancements like chemotherapy and IVF also *temporarily* lower GDP per capita; yet in the long run, they typically produce positive results. Near-term advancements in aging science that extend lifespan in imperfect health follow a similar trajectory, resulting in overwhelmingly positive net present values in the long run.

The role of what we call “first-generation” trials (like TAME) in collecting and validating biomarkers of aging and endpoints cannot be overstated. These trials will likely be essential to legitimize the possibility of running clinical trials with biological aging as an endpoint. Yet in the longer run, medicines will likely be developed to *prevent* the burdens of age-related disease and social care in still-healthy, working-age adults. While overweight or older adults may benefit from existing drugs, healthy or younger adults may not. What we call “second-generation therapeutics” could be developed to delay biological aging in still-healthy, working-age adults, with the same safety and efficacy profile now reliably achieved for sick, overweight, often retired adults. Many market failures stand in the way of such a preventative therapeutic class — but it can be developed. And as we demonstrate in Table 2 of this manuscript, the social and economic returns (in terms of lives extended and GDP growth) from such a therapeutic would be far higher.

Lastly, and counterintuitively, our results suggest the effects of slowing brain aging by 1 year on GDP per capita are higher than slowing *overall* biological aging by 1 year. This is because improvements in non-cognitive functions have diminishing returns, and a full one-year shift backwards in mortality rates (which a one-year shift in overall aging affords) translates into more chronologically older adults alive at any given time, many of whom will be eligible for Social Security. As well, the returns from improving the age of other organs (e.g. the kidneys, or the ovaries) are not immediate, and sometimes even *reduce* GDP temporarily—for instance, by increasing birth rates. Lastly, the same is true of our simulation on first-generation (existing, FDA-approved) therapeutics repurposed for biological aging: because they impact older adults over the age of 65 and with one pre-existing disease of aging, they consist of *treatment* more than *prevention*, and therefore do not offer positive impacts on GDP per capita. We repeat that while this result remains welcome in the short run (older adults with existing diseases of aging deserve to be treated), the long-term potential of aging science—which we simulate in our second-generation therapeutic—is to *prevent* age-related decline in still-healthy adults.

Another reason we focus on aggregate quantities is because so much of the returns from our simulations come from people who would not otherwise be alive. Increases in fertility rates create new humans; decreases in mortality rates afford existing humans with longer life. Both result in more life years in aggregate in our counterfactual scenarios. Extending healthspan results in more people years, and with our simulations, we have aimed to fully value those.

### Interviews with 102 scientists

We interviewed 102 scientists for this project. Geographically, roughly half of all scientists we interviewed were based in the vicinity of either Boston or San Francisco. Our interviewees represent a diverse and influential cross-section of the global biotechnology ecosystem. They include leading scientists, physicians, biotechnology founders, and investors from institutions ranging from Harvard, Stanford, and the Buck Institute for Research on Aging (three institutions which, together, host roughly half our interviewees) to Columbia, the University of Rochester, Birmingham, Cambridge, ETH Zurich, University of Toronto, Wake Forest, Yale, University of Texas Medical Branch, Scripps, German Center for Neurodegenerative Diseases, and not-for-profit foundations in the field, including The American Federation for Aging Research, the Methuselah Foundation, the Hevolution Foundation, the XPRIZE Foundation, and the Amaranth Foundation. 97 of the scientists completed an online survey, and 8 of them were interviewed in person, over email, or over a video call. We provide details on the completed surveys with 97 respondents below.

We note the purpose of this survey is conceptual rather than quantitative. Aging biology is a rapidly evolving field with no agreed-upon probability-of-success literature. It has not been our aim to fill this gap. Our surveys document scientific consensus on what constitutes plausible near-term advancements in aging science, but we do not assume that today’s scientists can reliably predict the future. This is why, though we document scientists’ views on timelines and investment amounts required for progress in aging science, we do not use these values to inform our own modeling approach. Our simulations focus on small, empirically grounded year-shifts (e.g., slowing ovarian aging by one year), and our surveys justify why such increments are realistic and relevant to policy discussions. In other words, our ambition is for this survey to help *contextualize* the economic results we present in this paper.

##### Select data findings from our surveys

1. **59.4%** of respondents believe that slowing aging by 5 years is already possible *today* with low-tech solutions like exercise and/or existing therapeutics. **31.3%** of respondents believe it will be possible in 10–20 years, while **9.4%** believe it will likely take 20–40 years. **No scientist** replied “it is impossible to safely delay biological aging.”
2. **38.3%** of respondents believe we will be able to slow biological aging by 30 years within 10–20 years. **27.7%** responded “this will likely be possible in 20–40 years,” and **30.9%** that “this will likely be possible before the end of this century.” **3.2%** replied it is impossible to safely delay biological aging by 30 years.
3. **37.2%** of respondents believe there is no ceiling to how much we can extend healthy lifespan. **54.3%** believe that if there is a ceiling, it is far higher than the current ∼120-year maximum lifespan. **8.5%** replied there *is* a ceiling, and “even if we compress morbidity, humans will still inevitably die at age 120 at most.” This latter response highlights how different scientists understand aging: many biotechnologists we interviewed believe that truly improving how we age would necessarily increase our *maximum* lifespan, even if maximum lifespan is not an ideal endpoint for a trial, since it would involve several decades of research. In our model, all agents necessarily die at age 100, in part because there is little empirical data on how adults aged 100 and older behave in the economy. We leave it up to readers to decide if they believe we are simulating “slowing” or “reversing” biological age, depending on their own definition of aging.
4. Only **7.6%** of respondents believe it would take *more* than $50 billion in funding to arrive at a future where biological aging can be *reversed* by 5 years. (This is a simulation five times more ambitious than the most ambitious of our simulations, which considered a 1-year shift in biological aging.) **45.7%** of respondents believe this could be done with ∼$1 billion in funding, since funding *direction* is more important than funding *amount*. **46.7%** of respondents believe it would take an investment ranging between $1–$50 billion.
5. **63.2%** of respondents believe a combination of intracellular therapeutics and “replacement” approaches (e.g. replacing the extracellular matrix to improve brain aging) will likely be needed to target biological aging in the coming decade. **30.5%** of respondents believe human aging will be managed *mostly* via intracellular/epigenetic therapeutics. **6.3%** replied most of human aging will likely need to be treated via cell/tissue/organ replacement. **Zero** scientists replied that “biological aging cannot be therapeutically targeted.”

#### Survey responses

| Choices | Responses | Percentages |
| --- | --- | --- |
| This is possible today with low-tech solutions like exercise and/or existing therapeutics. | 57 | 59.4% |
| This will likely be possible in ~10-20 years. | 30 | 31.3% |
| This will likely be possible in ~20-40 years. | 9 | 9.4% |
| It's impossible to safely delay biological aging. | 0 | 0% |

By when do you believe we'll be able to slow overall biological aging by 30 years? 94 out of 96 people answered this question.
| Choices | Responses | Percentages |
| --- | --- | --- |
| This will likely be possible in ~10-20 years. | 36 | 38.3% |
| This will likely be possible in ~20-40 years. | 26 | 27.7% |
| This will likely be possible before the end of this century. | 29 | 30.9% |
| It's impossible to safely delay biological aging by 30 years. | 3 | 3.2% |

3 Is there a ceiling to how much we can extend healthy lifespan? 94 out of 96 people answered this question.
| Choices | Responses | Percentages |
| --- | --- | --- |
| No ceiling. | 35 | 37.2% |
| Maybe, but the ceiling is far higher than the record ~120 years old. | 51 | 54.3% |
| Yes, even if we compress morbidity, humans will still inevitably die at age ~120 at most. | 8 | 8.5% |

4 How much funding will it take to engineer a future where biological aging is reversible by 5 years? 92 out of 96 people answered this question.
| Choices | Responses | Percentages |
| --- | --- | --- |
| It could be done in this decade with less than \$1B. Funding *direction* is more important than funding *amount*. | 42 | 45.7% |
| This will likely require more than \$1B in funding, and a 10-20 year timeline. | 43 | 46.7% |
| This will likely require more funding than the entire NIH budget, at \$50B, over several decades. | 7 | 7.6% |
| It will never be done. | 0 | 0% |

5 In your view, how will biological aging most likely be treated? 95 out of 96 people answered this question.
| Choices | Responses | Percentages |
| --- | --- | --- |
| Most of human aging will be managed via intracellular/epigenetic therapeutics. | 29 | 30.5% |
| Most of human aging will need to be treated via tissue/cell/organ replacement (e.g. replacing the extracellular matrix to improve brain aging). | 6 | 6.3% |
| There isn't enough evidence yet for either A or B. We'll continue to need a combination of therapeutics & interventions in the coming decade. | 60 | 63.2% |
| Biological aging cannot be therapeutically targeted. | 0 | 0% |

#### Mapping the TAME trial endpoints

The TAME trial’s primary outcome is a 20% reduction in outcomes including mortality and age-related chronic diseases (e.g. dementia, or stroke). The trial’s secondary outcome considers a 30% prevention in the decline of several biomarkers of aging (e.g. walking speed, bodily pain, cognitive function). The grouping of outcomes in TAME (mortality alongside chronic diseases) makes it difficult to parse out exact effects on mortality alone, making our mapping of these effects imperfect. To arrive at plausible shifts in mortality, we consider evidence across several clinical trials of metformin to date where all-cause mortality has already been rigorously measured. Diabetics on metformin have been documented to have an average mortality hazard rate 7% lower than age-matched healthy non-diabetics.^44^ This translates to a 6.1% reduction in mortality rates, and it is the (conservative) number we use to approximate the expected effects on mortality of a first-generation therapeutic.

The precise effects of existing therapeutics on mortality are difficult to quantify precisely. Baseline mortality rates in diabetics are higher than in non-diabetics, and this suggests the possibility of a stronger effect on all-cause mortality than 6.1%. One metastudy, considering over *one million* research participants throughout several studies, suggests a reduction in all-cause mortality of 33% given metformin use in patients with coronary artery disease.^45^ We chose to apply the far more conservative estimate of 6.1% reduction in all-cause mortality, in part because other studies suggest metformin’s effects on mortality may be confounded by biases and selective reporting.

We begin our simulations with the most conservative and well-documented numbers available. Starting with the endpoint of 6.1% decline in all-cause mortality, we mapped the first phase of TAME to estimate who might benefit from this intervention. The trial designers (whom we interviewed) estimate that roughly 50% of non-diabetic adults over the age of 65 could benefit from metformin. Yet actual enrollment in the trial is expected to total 11% of adults over age 65. This is because a) many people will decline to adhere to metformin over the trial duration of 6 years; b) some will be disqualified due to severe preexisting conditions (e.g. late-stage cancer); c) some will not have exactly one pre-existing disease of aging, as the trial demands; and d) the TAME trial disqualifies participants with an existing diabetes diagnosis, since its goal is to measure the effects of an existing therapeutic on biological aging. This actual enrollment rate of 11% is a conservative estimate of the treatment population. Yet it serves as a useful benchmark for the added economic effects of metformin use by the normally aging, non-diabetic population. 11% is the baseline number we use.

From there, we arrive at a population-wide reduction in mortality of 0.67%, which can be considered the “intent to treat effect” across the population, of which 11% adopt the treatment. The 0.67% intent to treat effect is arrived at from the product of the treatment effect of the drug, a 6.1% decline in all cause mortality, and the 11% share of the population aged 65+ whom we assume adopt this treatment (i.e., 0.67 = 0.061 * 0.11). The endpoints of the TAME trial, mapped in this way, result in a 0.67 percent decline in mortality rates *on average* for all US adults aged 65 and older. This percentage shift in population-wide mortality, again, is far lower than what previous studies on the effects of metformin on mortality suggest. This is because we consider a subset of the US population of older adults who a) is *not* diabetic, b) *is* suffering from exactly one disease of aging, and c) would adhere to metformin use over the trial’s duration.

It may be the case that even if only 11% of older adults would actually enroll in the TAME trial, once the trial is complete — and if it successfully proves that metformin has beneficial, pleiotropic effects on the human healthspan — a larger percentage of older, non-diabetic adults would adhere to metformin. This is why we *also* simulate how existing therapeutics could be favorable for up to 50% of all non-diabetic adults aged 65 and older. The CDC estimates 74% of this age group are *not* diabetic, and the trial designers expect that half this population (37%) would be eligible. Assuming, further, a medication adherence rate of 50% (roughly that of statins), we would arrive at an added beneficial impact of metformin on 18.5% of all US adults aged 65 and older. Below, we present this result as an alternative to the lower bound (11%).

We translate the TAME trial’s 30% prevention in decline of function into commensurate effects on productivity by finding the decline in labor productivity among 65-80 year olds (the age group of trial participants) measured by hourly earnings. For US workers, the average decline in hourly earnings from age 65 to 80 is about 66%. We infer from the work of Lindqvist and Vestman (2011) that the functions measured in the TAME trial — which are predominantly physical, but also include cognitive function — account for about 30% of labor productivity. We then adjust for sample selection (just 11% of the population of adults aged 65 and older is impacted), and arrive at a marginal increase in labor productivity among those aged 65 and older of 0.33% as the product of **30% x 66% x 30% x 11%**.

We acknowledge that the mapping of function as productivity is imperfect at best. Yet given the expected outcome of the TAME trial in additionally lowering the incidence rates of six specific diseases (Incident Myocardial Infarction, stroke, Congestive Heart Failure, Mild Cognitive Impairment, dementia, cancer) — which we do not compute in our simulation of mortality rates — the assumption of 30% prevention in decline of function appears just. In reality, the effects of this first-generation drug on mortality and productivity are unlikely to be as neatly divorced as they seem in our model. We have attempted to map these effects as faithfully as possible, while acknowledging that this mapping will have been imprecise.

Caloric restriction for overweight older adults has already proven to offer a 15% reduction in all-cause mortality, and to meaningfully reduce disease rates.^46^ Weight-loss and diabetes drugs like semaglutide (i.e., Ozempic) and metformin may incidentally target the biology of aging by tweaking mTOR (a protein that helps control cell division and survival) and insulin signaling. Calorie-restriction mechanisms *may* even marginally slow aging in non-obese, normally aging humans. Yet no trials have been completed in humans, and existing drugs like metformin have shown undesirable side effects in young adults.^47^ It is likely that in the category of existing therapeutics, GLP-1 a2gonists or rapamycin would have superior effects on mortality, but they may or may not have a less favorable safety profile, and their effects on the biology of aging are unknown. Given a lack of clinical trials for the latter drugs in particular, these assumptions remain speculative. We consider the TAME trial because the effects of metformin are particularly well-documented *on older adults*, showing an often favorable safety profile.

## Data Availability

All data are open-source, publicly available, and from public sources. Sources are documented in the text and all data used to create figures in the paper are available at: https://github.com/OpenSourceEcon/MacroAgingNature

More information on each parameter of the model we use can be found below: https://pslmodels.github.io/OG-USA

The typeform survey with the 7 questions posed to 102 scientists can be found below: https://form.typeform.com/to/cP9HV05J

The list of responses to the surveys can be found below: https://form.typeform.com/report/cP9HV05J/O3lchbyjLRQfKogW

Below, readers can also inspect model parameters and simulation results in a friendlier format for users not trained in economics: https://profound-brioche-f9933d.netlify.app/

## Code Availability

All code necessary to run model simulations and create figures and tables for this article are available online in the GitHub repository https://github.com/OpenSourceEcon/MacroAgingNature.

## Acknowledgments

We thank George Church, Tyler Cowen, Eric Budish, Richard Freeman and three anonymous referees for their thoughtful comments on this paper.

## Supplemental Information

Baseline simulations with optimistic and pessimistic scenarios

**Supplementary Information Table 1.** OG-USA simulation results for various medical breakthroughs in aging therapies, optimistic and pessimistic scenarios.

| Sim # | Simulation Description | Mortality decrease in years | Productivity increase in years | Fertility increase in years | Annual change in US GDP (Avg. over 2045-2064) | NPV of increased GDP over several decades | Increase in 2050 population |
| --- | --- | --- | --- | --- | --- | --- | --- |
| 1 | **Slowing brain aging by 1 year** | **0.20** | **0.70** | **0.0** | **\$201B** | **\$8.9T** | **268K** |
| 1a | Pessimistic version of simulation 1 | 0.16 | 0.56 | 0.0 | \$157B | \$7.1T | 214K |
| 1b | Optimistic version of simulation 1 | 0.24 | 0.84 | 0.0 | \$246B | \$10.7T | 322K |
| 2 | **Slowing reproductive aging by 1 year** | **0.02** | **0.10** | **1.0** | **\$9B** | **\$9.3T** | **391K** |
| 2a | Pessimistic version of simulation 2 | 0.016 | 0.08 | 0.8 | \$4B | \$7.4T | 313K |
| 2b | Optimistic version of simulation 2 | 0.024 | 0.12 | 1.2 | \$15B | \$11.1T | 470K |
| 3 | **Slowing overall aging by 1 year for all adults age 40+** | **1.0** | **1.0** | **1.0** | **\$408B** | **\$27.1T** | **1.7M** |
| 3a | Pessimistic version of simulation 3 | 0.8 | 0.8 | 0.8 | \$321B | \$21.6T | 1.4M |
| 3b | Optimistic version of simulation 3 | 1.2 | 1.2 | 1.2 | \$496B | \$32.7T | 2.1M |
| 4 | **Slowing overall aging by 1 year for all adults age 65+** | **1.0** | **1.0** | **1.0** | **\$326B** | **\$14.3T** | **1.2M** |
| 4a | Pessimistic version of simulation 4 | 0.8 | 0.8 | 0.8 | \$256B | \$11.4T | 939K |
| 4b | Optimistic version of simulation 4 | 1.2 | 1.2 | 1.2 | \$397B | \$17.4T | 1.4M |

**Supplementary Information Table 2.** OG-USA simulation results for advancements in our ability to *measure* biological aging, optimistic and pessimistic scenario.

| Sim # | Simulation Description | Mortality decrease in years | Productivity increase in years | Fertility increase in years | Annual change in US GDP (Avg. over 2045-2064) | NPV of increased GDP over several decades | Increase in 2050 population |
| --- | --- | --- | --- | --- | --- | --- | --- |
| 5 | **1st generation, all adults ages 65+, 1 disease of aging, 11% of non-diabetic adults** | **0.82** | **0.73** | **0.0** | **\$40B** | **\$2.4T** | **275K** |
| 5a | Pessimistic version of simulation 5 | 0.66 | 0.58 | 0.0 | \$39B | \$2.3T | 260K |
| 5b | Optimistic version of simulation 5 | 0.98 | 0.88 | 0.0 | \$42B | \$2.5T | 294K |
| 6 | **1st generation, all adults ages 65+, 18.5% of non-diabetic adults** | **0.81** | **0.72** | **0.0** | **\$79B** | **\$4.2T** | **460K** |
| 6a | Pessimistic version of simulation 6 | 0.65 | 0.58 | 0.0 | \$76B | \$4.0T | 435K |
| 6b | Optimistic version of simulation 6 | 0.97 | 0.86 | 0.0 | \$82B | \$4.3T | 491K |
| 7 | **2nd generation, all adults ages 40+, 50% of non-diabetic adults** | **0.82** | **1.44** | **0.0** | **\$505B** | **\$22.2T** | **1.1M** |
| 7a | Pessimistic version of simulation 7 | 0.66 | 1.15 | 0.0 | \$498B | \$21.9T | 1.1M |
| 7b | Optimistic version of simulation 7 | 0.98 | 1.73 | 0.0 | \$513B | \$22.7T | 1.2M |
| 8 | 2nd generation with organs on chips, all adults ages 40+, 50% of non-diabetic adults, 20% faster development | 0.82 | 1.44 | 0.0 | \$522B | \$23.2T | 1.3M |
| 8a | Pessimistic version of simulation 8 | 0.66 | 1.15 | 0.0 | \$515B | \$22.9T | 1.3M |
| 8b | Optimistic version of simulation 8 | 0.98 | 1.73 | 0.0 | \$532B | \$23.7T | 1.4M |

## Notes

https://github.com/OpenSourceEcon/MacroAgingNature

### Competing Interest Statement

The authors have declared no competing interest.

